# Heritability and family-based GWAS analyses of the *N*-acyl ethanolamine and ceramide plasma lipidome

**DOI:** 10.1101/815654

**Authors:** Kathryn A. McGurk, Simon G. Williams, Hui Guo, Hugh Watkins, Martin Farrall, Heather J. Cordell, Anna Nicolaou, Bernard D. Keavney

## Abstract

Signalling lipids of the *N*-acyl ethanolamine (NAE) and ceramide (CER) classes are emerging as novel cardiovascular disease biomarkers. We sought to establish the heritability of plasma NAEs (including the endocannabinoid anandamide) and CERs, and identify common DNA variants influencing the circulating concentrations of the heritable lipid species. Nine NAE and sixteen CER species were analysed in plasma samples from 999 members of 196 British Caucasian families, using targeted mass spectrometry (UPLC-MS/MS). Heritability was estimated and GWAS analyses were undertaken; all target lipids were significantly heritable (h^2^ = 36%-62%). A missense variant (rs324420) in the gene encoding the enzyme fatty acid amide hydrolase (*FAAH*), which degrades NAEs, associated at GWAS significance (P<2.15×10^−8^) with four NAEs (DHEA, PEA, LEA, VEA). The A allele of this SNP was associated with a 0.23 SD per-allele increase in plasma NAE species. Additionally, we found association between rs680379 in the *SPTLC3* gene, which encodes a subunit of the rate limiting enzyme in CER biosynthesis, and a range of CER species (e.g. CER[N(24)S(19)]; P =4.82×10^−27^). We also observed three novel associations (*CD83, SGPP1, FBXO28-DEGS1*) influencing plasma CER traits, two of which (*SGPP1* and *DEGS1*) implicate CER species in haematological phenotypes. NAE and CER are substantially heritable bioactive lipids, influenced by SNPs in key metabolic enzymes.

## 1. Introduction

Genetic studies in large numbers of individuals have identified loci where common genetic variation influences the prevalence of the major plasma lipid species, such as HDL- and LDL-cholesterol, and triglycerides (1,2). Recent advances in targeted lipidomics have enabled quantitative analyses of a greater proportion of the mediator lipidome in blood, supporting attempts to identify genetic associations for low-concentration bioactive lipid species to potentially find cardiovascular disease biomarkers.

Bioactive lipids of the *N*-acyl ethanolamine (NAE) and ceramide (CER) classes have potent roles in inflammation and immunity (3–5). NAEs are fatty acid derivatives, derived from membrane phospholipids and degraded by the enzyme fatty acid amide hydrolase (*FAAH*; Figure 1). This class of bioactive lipids includes the endocannabinoid anandamide (AEA), the nuclear factor agonist palmitoyl ethanolamide (PEA), and a number of other species with roles in neuronal signalling, pain and obesity (6–9). The contribution of genetic factors to the blood levels of NAEs has only been studied for one NAE species, oleoyl ethanolamide (OEA), in a study using untargeted mass spectrometry (10).

**Figure 1:**
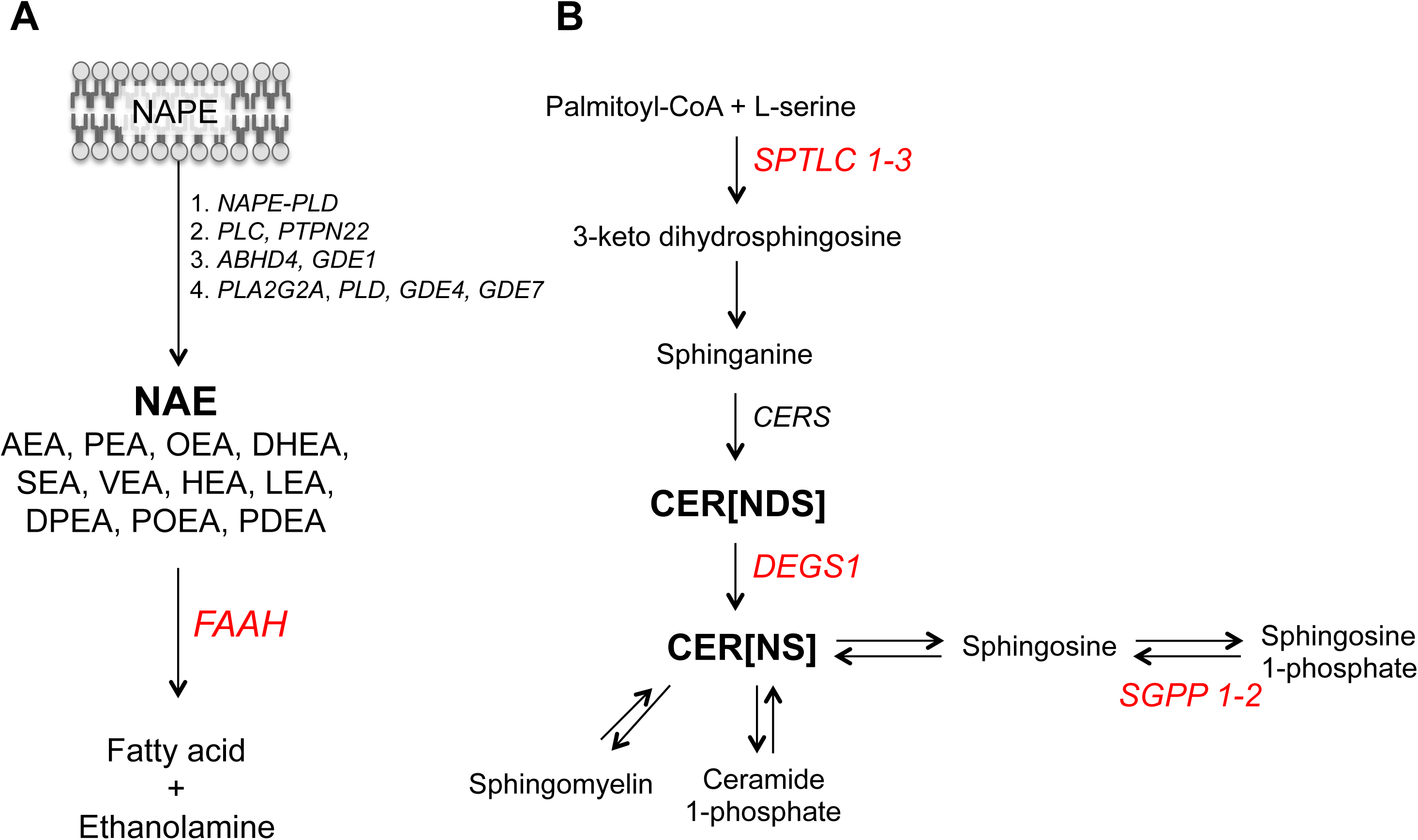
Schematic overview of the biosynthetic pathways for (A) *N*-acyl ethanolamines and (B) ceramides. A) *N*-acyl ethanolamine (NAE) species, including the endocannabinoid anandamide (AEA), are produced through four independent enzymatic pathways from the membrane phospholipid precursor (*N*-acyl phosphatidylethanolamine; NAPE). Fatty acid amide hydrolase (*FAAH*) degrades NAEs to free fatty acids (such as arachidonic acid for AEA) and ethanolamine. B) Ceramide (CER) species are biosynthesised via the enzyme serine palmitoyltransferase (*SPTLC 1-* 3) that converts palmitoyl-CoA and L-serine to 3-keto dihydrosphingosine, in the rate-limiting step of the sphingolipid *de novo* pathway. The resulting dihydrosphingosine is coupled to various fatty acids via ceramide synthases (*CERS*) to generate dihydroceramides CER[NDS] that are further converted to CER[NS] via the enzyme delta 4-desaturase (*DEGS1*). Conversion of these pro-apoptotic CER[NS] species to sphingosine and sphingosine 1-phosphate, with roles in cell survival, degrades ceramides through reversible reactions. CER[NS] are also reversibly converted to sphingomyelin or further metabolised to ceramide 1-phosphate). Measured lipid species are in bold; genes encoding enzymes are in italics; genes identified through SNPs that associated at GWAS with circulating lipid levels are in red.

Ceramides are members of the sphingolipid class, being derivatives of sphingoid bases (e.g. sphingosine and dihydrosphingosine) and fatty acids (Figure 1). The first and rate limiting step (11) of their *de novo* biosynthesis is catalysed by the enzyme serine palmitoyltransferase, a heterodimeric protein whose monomers are encoded by the *SPTLC1-3* genes. Ceramides play important roles in apoptosis (12). Recently, some circulating ceramide derivatives of 18-carbon sphingosine and non-hydroxy fatty acids (e.g. CER[N(16)S(18)]) have been suggested as novel biomarkers of cardiovascular death (13), type-2 diabetes, and insulin resistance (14). Furthermore, the contribution of genetic factors to these ceramides species has also been investigated (15–18).

In this study we analysed plasma NAEs and ceramides by mass spectrometry-based targeted quantitative lipidomics in 196 British Caucasian families comprising 999 individuals, and then investigated the heritability and common genetic variant associations. We show that these bioactive lipid mediators are substantially heritable, and demonstrate that plasma NAEs and a wide range of ceramides are influenced by SNPs in key metabolic enzymes (*FAAH, SPTLC3, DEGS1, SGPP1)*. Furthermore, we identify a novel inflammatory locus (*CD83*) associated with ceramide species, and implicate this group of bioactive lipid mediators in haematological phenotypes through Mendelian randomisation, using SNPs in *DEGS1* and *SGPP1* as instruments.

## 2. Results

### 2.1 Population characteristics

Plasma samples of 999 participants from 196 British Caucasian families were included in the genetic analyses. The families consisted of 1-24 members (mean of 5 members) with plasma available for lipidomics analyses (Figure S2). Participant descriptions are listed in Table 1.

**Table 1:** Summary statistics for the study participants. Data is shown as mean and standard deviation (SD) unless otherwise indicated; BMI, body mass index; WHR, waist-hip ratio.

| Trait | Mean (SD) |
| --- | --- |
| Gender | 47% Male |
| Hypertensive | 33% |
| Mean Blood Pressure | 138/83 mmHg |
| Age (years) | 49 (15) |
| BMI | 26.04 (4.33) |
| WHR | 0.86 (0.09) |
| Cholesterol (mmol/L) | 5.61 (1.20) |

### 2.2 Lipidomics descriptive statistics

Of the nine NAE species identified in plasma, palmitoyl ethanolamide (PEA) was at highest abundance (1.89 ± 1.36 ng/ml [mean ± SD]), and of the sixteen plasma ceramide species, CER[N(24)S(18)] was most abundant (2.72 ± 1.29 nmol/ml), similar to previous studies (19–23).

### 2.3 Signalling lipid species are highly heritable

The signalling lipid species studied in this project were estimated to be substantially heritable (Figure 2, Table S8). The NAE species had estimated heritabilities ranging from 45% to 56% (P_adj_<4.28×10^−15^), with *N-*docosahexaenoylethanolamine (DHEA) producing the highest estimated heritability. The ceramide species had estimated heritabilities ranging from 36% to 62% (P_adj_<4.40×10^−13^), with CER[N(25)S(20)] and CER[N(24)S(16)] producing the highest estimated heritability.

**Figure 2:**
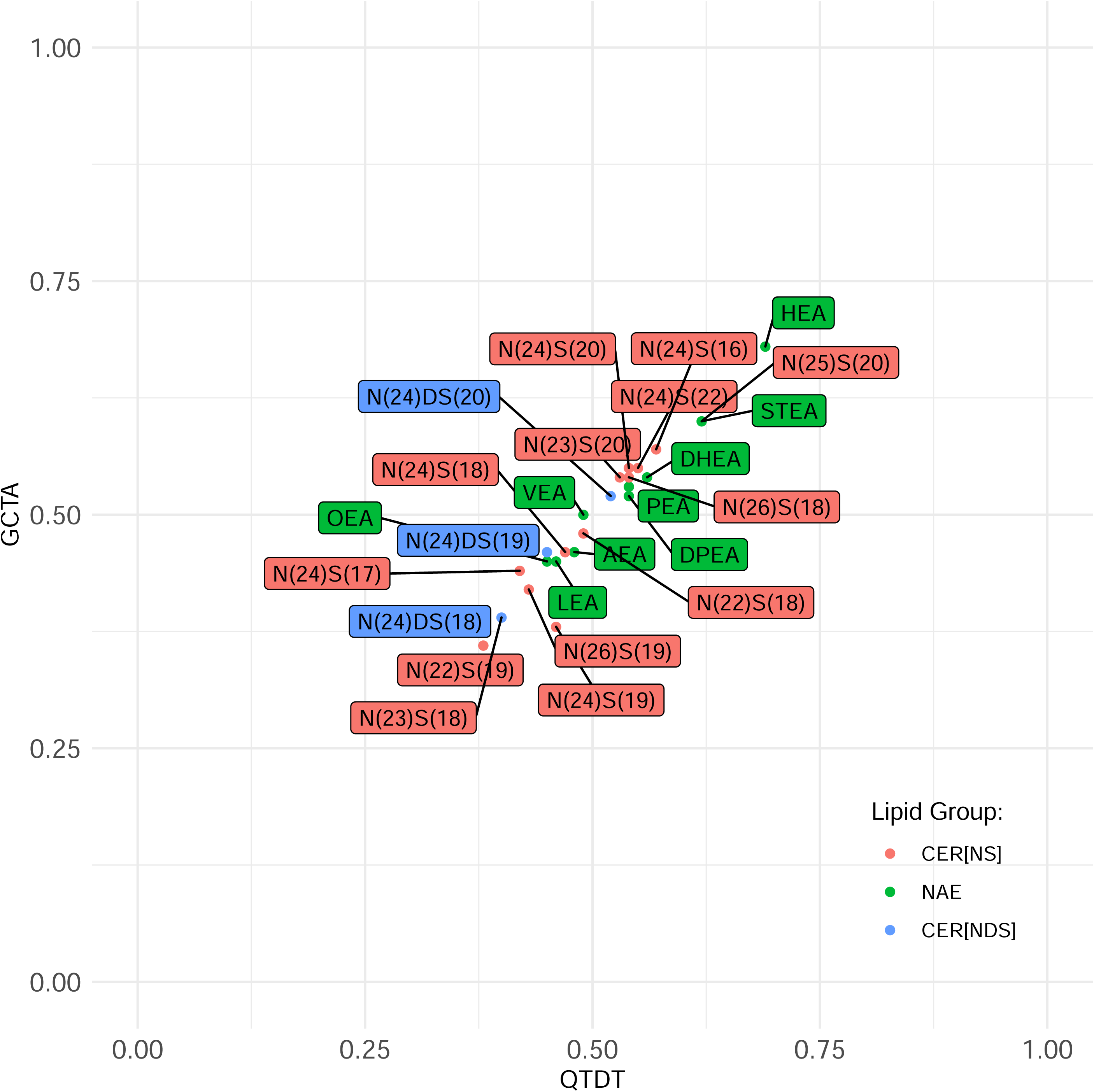
Heritability estimates of *N*-acyl ethanolamines and ceramides found in human plasma. This figure depicts the heritability estimated for each of 25 lipid species in 999 plasma samples using SNP-based GCTA software (y-axis) and reported pedigree-based QTDT software (x-axis). This data is presented in detail in Table S8.

### 2.4 Genome-wide association study of N-acyl ethanolamines

There were conventionally GWAS significant (P<5×10^−8^) associations between four NAEs (*N*-docosahexaenoyl ethanolamide, DHEA; *N*-linoleoyl ethanolamide, LEA; *N*-palmitoyl ethanolamide PEA; *N*-vaccinoyl ethanolamide, VEA), as well as the sum of all NAEs (sumNAE), with SNPs in the gene encoding fatty acid amide hydrolase (*FAAH*), which catalyses the degradation of NAEs (Figure 1, Table S9, Figure S3). The leading SNP is a missense variant (rs324420; C385A; P129T) and eQTL of *FAAH* in multiple tissues including whole blood. Presence of the missense variant causes the enzyme to display normal catalytic properties but decreased cellular stability (24) by enhanced sensitivity of the enzyme to proteolytic degradation (19). The magnitude of the genetic effect was considerable (Figure 3); the A allele of the lead SNP rs324420 caused a 0.23 SD per-allele increase in the plasma NAE species.

**Figure 3:**
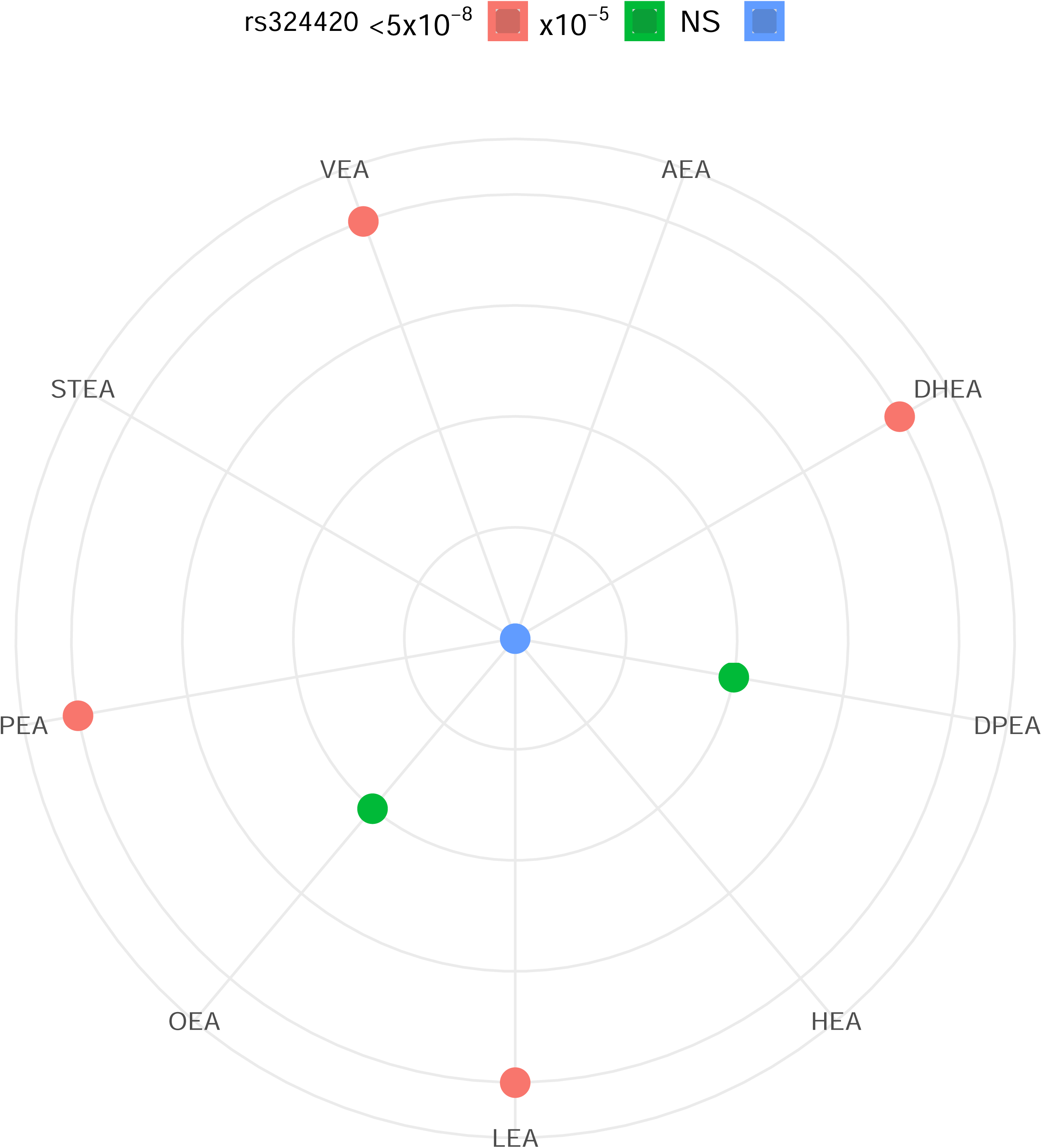
Family-based GWAS results for *N*-acyl ethanolamines and the lead SNP in fatty acid amide hydrolase (*FAAH*). A) The radar plot depicts the P-value for association between the lead SNP and eQTL of *FAAH* (rs324420) and each of 9 NAE species in 999 plasma samples. The P-values were grouped into “<5×10^−8^” (P<5×10^−8^, outermost ring), “x10^−6^” (P=5.0×10^−8^ - 9.9×10^−6^ [of which there are no NAE species]), “x10^−5^” (1.0×10^−5^ - 9.9×10^−5^), and “NS” (not significant) at the center of the radar. B) Trend in concentrations of plasma NAE species separated by FAAH rs324420 genotype. The figure depicts the mean standardised residuals of the NAE species in participants with the three genotypes at rs324420. The mean is shown with standard error. 51 participants had the AA genotype, 310 had the AC genotype, and 638 had the CC genotype in the cohort. C) LocusZoom plot of the association of PEA with *FAAH* SNP rs324420. The LocusZoom plot depicts the association of *N*-acyl ethanolamine lipid species PEA with *FAAH* SNP rs324420 on chromosome 1 in 993 plasma samples. The r^2^ for each SNP is depicted in colour. The plot was created using the LocusZoom plot tools at http://locuszoom.sph.umich.edu/.

### 2.5 Genome-wide association study of ceramides and related sphingolipids

Seven CER[NS] and two CER[NDS] species were significantly associated with SNPs in an intergenic region on chromosome 20 (Figure 4, with example Manhattan plot depicted in Figure S4, and further details in Table S11). Assessing the SNPs using GTEx confirmed them as liver eQTLs (Table S11) found 20,000 bases downstream of the gene encoding the third subunit of serine palmitoyltransferase (*SPTLC3*; Figure 6), which catalyses the rate-limiting step (11) of CER biosynthesis (Figure 1). The SNPs are associated with differences in the expression of the *SPTLC3* gene in the liver. Associated SNPs had considerable phenotypic effects, for example the A allele of the SNP rs680379 was associated with a 0.30 SD per-allele increase in plasma ceramides. Furthermore, the summed total of all ceramide species with 24-carbon non-hydroxy fatty acids, and, independently, those with 19- and 20-carbon sphingosine bases, were found associated with the same SNPs at the *SPTLC3* locus (Table S11). A novel association was identified for CER[N(26)S(19)] at a locus on chromosome 6, upstream to the gene for inflammatory protein CD83 (e.g. rs6940658, P=2.07×10^−8^; depicted in Figure S5, with further details in Table S11).

**Figure 4:**
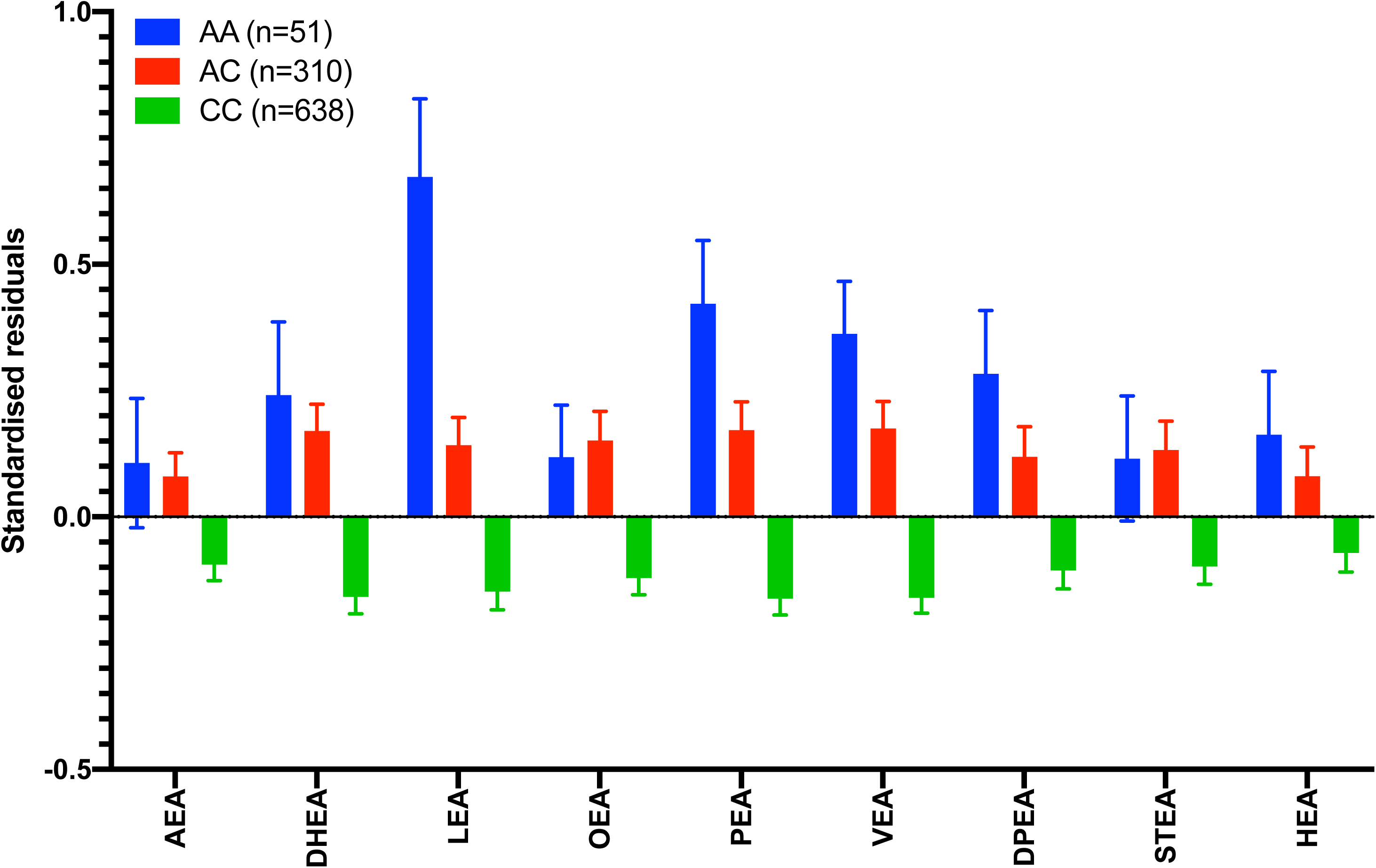
Family-based GWAS results for CER[NS] and precursor CER[NDS] with an exemplar SNP in serine palmitoyltransferase (*SPTLC3*). A) The radar plot depicts the P-value for association between the lead SNP and liver eQTL of *SPTLC3* (rs680379) with the 13 CER[NS] and 3 CER[NDS] species in 999 plasma samples. The P-values were grouped into “<5×10^−8^” (P<5×10^−8^, outermost ring), “x10^−6^” (P=5.0×10^−8^ - 9.9×10^−6^, “x10^−5^” (1.0×10^−5^ - 9.9×10^−5^), and “NS” (not significant) at the center of the radar. B) Trend in concentrations of plasma ceramide species separated by *SPTLC3* rs680379 genotype. The figure depicts the mean standardised residuals of the ceramide species in participants with the three genotypes at rs680379. The mean is shown with standard error. 409 participants had the GG genotype, 442 had the AG genotype, and 148 had the AA genotype in the cohort. C) LocusZoom plot of the association of CER[N(24)S(19)] with *SPTLC3* SNP rs680379 The LocusZoom plot depicts the association of CER[N(24)S(19)] with *SPTLC3* SNP rs680379 on chromosome 20 in 991 plasma samples. While there is a group of lead SNPs, this SNP was depicted as it has been identified previously to associate at GWAS with sphingolipid species. The r^2^ for each SNP is depicted in colour. The plot was created using the LocusZoom plot tools at http://locuszoom.sph.umich.edu/.

### 2.6 Association of ceramides and related traits with hematological phenotypes

The Gene Atlas Browser of PheWAS in the UK Biobank study was used to assess the association of significant SNPs identified here with the extensive number of phenotypes measured for the UK Biobank cohort. The ratio of CER[N(24)S(19)] to its biochemical precursor CER[N(24)DS(19)], is indicative of delta 4-desaturase, sphingolipid 1 (*DEGS1*) activity (Figure 1). A set of SNPs in the upstream region of the *DEGS1* gene on chromosome 1 associated with this ratio (P=4.34×10^−8^; Figure S6, Table S10). All significant SNPs were confirmed eQTLs of *DEGS1* in whole blood (Table S11). This locus associated with numerous blood cell phenotypes in the UKBiobank data (e.g. rs4653568 and mean platelet volume; P=4.77×10^−12^; Table S11).

The GWAS results for species CER[N(24)S(16)] showed an association with the *SPTLC3* region and also with a further set of SNPs upstream of the gene encoding sphingosine-1 phosphate phosphatase (*SGPP1*) (e.g. rs7160525, P=5.67×10^−10^; Figure S7). This enzyme is involved in the recycling of CER[NS] species from sphingosine and sphingosine-1-phosphate (Figure 1). All significant SNPs at this locus were also associated with blood cell phenotypes, identified in the UK Biobank data (e.g. rs7160525 and mean platelet volume, P=3.28×10^−29^; Table S11).

The significant SNPs identified at *SGPP1, DEGS1*, and *SPTLC3* genomic loci were used as instruments for two-sample Mendelian randomisation analyses (2SMR) using published blood cell count GWAS as outcome variables (25). The SNPs in *SGPP1* that associated with CER[N(24)S(16)] were significant (P_adj_<0.05) in influencing platelet, red blood cell, and white blood cell traits (Table S12). 2SMR analyses using the significant SNPs identified for *SPTLC3, FAAH, SGPP1, and DEGS1* as genetic instruments, did not suggest a causal role of the lipid species for which we detected GWAS significant association, in CVD (26). Neither did we find evidence for a causal association of the ceramide species measured here with type 2 diabetes using associated SNPs in *SPTLC3* (27).

## 3. Discussion

We show substantial heritability estimated for an array of signalling lipid mediators found in plasma and we identify GWAS significant associations between lipid species and DNA variants of the enzymes in their respective metabolic pathways. We have provided the first GWAS significant evidence of association between SNPs in the *FAAH* gene and four plasma NAEs (DHEA, LEA, PEA, and VEA). Additionally, we have extended the previously described association between SNPs in the *SPTLC3* gene and plasma ceramides to a wider range of species. Our results indicate that these two genes are the major loci influencing plasma levels of NAEs and CERs, respectively. In addition, we have shown novel SNP associations (*CD83, SGPP1, DEGS1)* influencing plasma ceramides species, that implicate ceramides in haematological phenotypes.

The NAE species DHEA, LEA, PEA, and VEA, associated with the SNP rs680379, a missense change in the NAE degradation enzyme FAAH. The association with PEA was identified previously in a single candidate gene study of mutations in *FAAH* in 114 subjects (28), which reported the same direction of effect on plasma AEA, PEA, and OEA species but with P-values insignificant at genome-wide levels (0.003<P<0.04). OEA is the only NAE species that has been previously associated with DNA variants at GWAS significance. An eQTL of *FAAH* (rs1571138, upstream to *FAAHP1*, P=5.15×10^−23^) that is in complete linkage disequilibrium with the lead SNP in our study, was identified in an untargeted study of blood lipids; OEA was the only NAE species measured in that study (10). Here, we found only a suggestive association with OEA (P=5.80×10^−5^), although we observed non-significant trends in the same direction for all NAE species with genotype at this SNP (Figure 3).

While the *FAAH* missense SNP rs324420 is not associated with any disease endpoints identified from GWAS to date, the A allele, associated with higher NAE levels, has been reported to increase the risk of polysubstance addiction and abuse [MIM: 606581] in three candidate gene studies totaling 863 cases and 2,170 controls (19,29,30). PheWAS analysis using the Gene Atlas UK Biobank online browser however, did not identify significant association in a similar number of UK Biobank cases of substance abuse/dependency (OR for A allele = 1.10; P = 0.14; 746 cases and 451,518 controls). It is possible that misclassification bias has affected the UK Biobank PheWAS; among the 451,518 UK Biobank participants assigned as controls, some reported dependencies on other substances and behaviours, such as coffee, cigarettes, prescription drugs, and gambling [UKBiobank data show case; http://biobank.ndph.ox.ac.uk/showcase/, accessed April 2019]. The potential implication of NAE species in addiction through the association with the *FAAH* SNP, warrants further investigation in larger numbers of cases.

As direct cannabinoid receptor 1 antagonist drugs have caused severe adverse psychiatric effects (31), FAAH inhibitors are being evaluated as an alternative approach to modulating endocannabinoid signalling (32,33). However, In 2016, a FAAH inhibitor resulted in severe neurological side-effects in a Phase I trial; this was hypothesised due to off-target drug effects rather than adverse on-target effects (34). As the functional *FAAH* SNP rs324420, which substantially impacts FAAH activity, did not associate with any adverse phenotypes in the UK Biobank, it is likely that on-target effects of FAAH inhibitors do not have substantial risks of causing conditions that occurred with appreciable frequency in UK Biobank.

Narrow-sense heritability has been estimated for ceramides in previous studies, showing estimated heritability of 9% to 51% (17,18). Here, we assess a different array of species and expand on these previous estimates to show that further CER[NS] and CER[NDS] are significantly heritable. The rs7157785 variant in sphingosine 1-phosphate phosphatase 1 (*SGPP1*), a ceramide metabolic enzyme, has been identified previously in GWAS of sphingomyelin (15,16,35), total cholesterol (36), glycerophospholipids (35), and the ratio of an unknown blood lipid (X-08402) to cholesterol (37). The novel association with CER[N(24)S(16)] we describe is consistent with the enzyme’s role in influencing CER[NS] production, through the formation of sphingosine (C18S) for CER[NS] biosynthesis (Figure 1). The other significant SNPs identified at the same locus and in linkage disequilibrium with the lead SNP, associated with this ceramide species and have been previously identified in further GWAS studies of blood phospholipids (38), red cell distribution width (25), sphingomyelin (38), and unknown blood metabolite X-10510 (37). All SNPs identified at this locus associated in the UKBiobank PheWAS assessment with multiple blood cell counts and other hematological phenotypes. Ceramides have been previously shown to stimulate erythrocyte formation through platelet activating factor (39). However, further studies will be required to identify the mechanism of the association between genetically determined plasma ceramide levels and blood cell phenotypes.

CER[N(26)S(19)] associated at GWAS significance with SNPs at a novel locus on chromosome 6, upstream to the gene encoding the inflammatory protein CD83 (P=2.07×10^−8^), a member of the immunoglobulin superfamily of membrane receptors expressed by antigen-presenting white blood cells, leukocytes, and dendritic cells (40). An interaction between CD83 and ceramides is currently unknown, but given the involvement of ceramide signalling in inflammation and immunity (41,42), it would be of interest to investigate further.

Association between some ceramide species and the *SPTLC3* SNP rs680379 has been identified previously through the use of shotgun lipidomics for four ceramide species (CER[N(22)S(18)], CER[N(23)S(18)], CER[N(24)S(18)], and CER[N(24:1)S(18)]) at GWAS significance (15,16). Here, we identify associations between an additional seven CER[NS] and two CER[NDS] plasma species and this SNP, and with other eQTLs of serine palmitoyltransferase at the same locus; as this enzyme is the rate limiting step for the *de novo* biosynthesis of ceramides, this association may have wider implications. While we did not find a signficiant association with all CER and *SPTLC3* at GWAS, we observed non-significant trends in the same direction for all ceramide species with genotype at the rs680379 SNP (Figure 4). The information gathered from the eQTL analysis highlights all of the *SPTLC3* confirmed eQTLs act in the liver, which is a major site for plasma ceramide biosynthesis. Neither PheWAS analysis in UK Biobank, nor 2SMR analysis, identified significant disease associations with the *SPTLC3* locus. A number of CER[NS] species have been studied as potential biomarkers of cardiovascular disease and diabetes (13,43), and data from others has suggested that the *SPTLC3* locus is associated with these ceramides (15,16). The extent to which specific species have a role in cardiovascular disease remains debated (44,45).

A limitation of this study is its size (999 participants), which limits the power to detect small effects; however, our study is the largest of which we are aware that has analysed this range of plasma NAE and ceramide species to date. The inclusions of non-fasting samples that are subjectively haemolysed or contain white blood cells may add noise to the analysis that would impair the detection of weaker genetic effects. However, this may not be a signficiant issue as a study on blood ceramide lipidomics analysis did not find differences between serum and plasma samples, and fasting and non-fasting samples (46).

The associations we have uncovered suggest that further investigation of heritable lipid species for which no GWAS association was found in this study, in larger cohorts and ethnically diverse populations, would be of interest.

## 4. Materials and Methods

### 4.1 Family recruitment

Families were recruited for a quantitative genetic study of hypertension and other cardiovascular risk factors, and selected via a proband with essential hypertension (secondary hypertension was excluded using standard clinical criteria) as previously described (47). Probands were recruited from outpatients attending the John Radcliffe Hospital, Oxford hypertension clinic, or via their family doctors. Included family members were U.K. residents of self-reported White European ancestry and were required to consist of 3 or more siblings quantitatively assessable for blood pressure if one parent of the sibship was available for blood sampling, or 4 or more siblings if no parent was available. The hypertensive proband could be either in the sibship or parental generation. First, second and third degree relatives were then recruited to assemble a series of extended families. The participants were fully phenotyped for blood pressure (using ambulatory monitoring), cardiovascular risk factors, blood biochemical measures, and anthropometric traits. Non-fasting blood samples were collected, plasma separated, and stored at -80°C until lipidomic extraction. DNA was extracted from whole blood by standard methods. The collection protocol obtained ethical clearance from the Central Oxford Research Ethics Committee (06/Q1605/113) and it corresponds with the principles of the Declaration of Helsinki. Written informed consent was obtained from all participants. This cohort of extended families has previously been shown to have adequate power to detect moderate-sized genetic influences on quantitative traits (48,49).

### 4.2 UPLC/ESI-MS/MS mediator lipidomics

Plasma samples were extracted and analysed by mass spectrometry as previously described (50–52). Briefly, lipids were extracted from plasma (1 mL) using chloroform-methanol in the presence of internal standards: CER[N(25)S(18)] (50 pmol/sample; Ceramide/Sphingoid Internal Standard Mixture I, Avanti Polar Lipids, USA) for CER, and AEA-*d*8 (20 ng/sample; Cayman Chemical Co., USA) for NAE. Targeted lipidomics was performed on a triple quadrupole mass spectrometer (Xevo TQS, Waters, UK) with an electrospray ionisation probe coupled to a UPLC pump (Acquity UPLC, Waters, UK). Ceramide species were separated on a C8 column (2.1 × 100 mm) and NAE were separated on a C18 column (2.1 × 50 mm) (both Acquity UPLC BEH, 1.7µm, Waters, UK). NAE species were quantified using calibration lines of synthetic standards (Cayman Chemical); relative quantitation of ceramides was based on the internal standard (Avanti Polar Lipids) (51,53). Thus, the concentration of plasma NAE species are reported in pg/ml, while the relative abundance of plasma ceramide species is reported in pmol/ml. Pooled plasma samples from healthy volunteers were used to create quality control samples that were extracted and analysed blindly alongside the familial samples. Detailed quality control information can be found in the Supplemental Methods. The CER[NS] notation (54) denotes a ceramide that contains a non-hydroxy fatty acid attached to a sphingosine base, for example, a 16-carbon non-hydroxy fatty acid joined to a 18-carbon sphingoid base is denoted as CER[N(24)S(18)], where N(24) represents a 24-carbon non-hydroxy fatty acid, and S(18) represents a 18-carbon sphingosine base attached. This species is also denoted as Cer(d18:1/24:0) in literature.

### 4.3 Statistical analysis

#### 4.3.1 Covariate adjustment

Systematic error was considered from a variety of sources and assessed for collinearity; mass spectrometry batch and a trait created to adjust for sample abnormality (haemolysis or presence of white blood cells; present in 14% of samples) were included as potential covariates. Ascertainment selection was modelled via binary hypertension status. The resulting concentrations for each lipid species from the pooled quality control samples were used for adjustment of systematic errors during extraction, quantitation, and data processing. The final set of potential covariates included mass spectrometry batch, sample abnormality, quality control sample measures, age, age^2^, sex, hypertension status, BMI, and total cholesterol. The lipid measurements were assessed for effect of potential covariates using stepwise multiple linear regression to identify the best set of predictors, using the ‘caret’ package and ‘leapSeq’ method in R (version 3.5.2) (see Table S4 for predictors). Multiple linear regression of the best predictors was undertaken using the ‘lm’ function in R. Residuals from the covariate-adjusted regression models were standardized to have a mean of 0 and a variance of 1. Outliers were assessed using the R package ‘car’, assessing each observation by testing them as a mean-shift outlier based on studentized residuals, to remove the most extreme observations (Bonferroni P-value of P<0.05). Missing values were coded as such in the genetics analyses. As lipid mediators can exert individual bioactivities, all lipid species were treated uniquely for all analyses, intra-class correlation analyses are depicted in Figure S1.

#### 4.3.2 Genome-wide genotyping quality control

Genotyping was performed using the Illumina 660W-Quad chip on 1,234 individuals (580 males and 654 females) including 248 founders, at 557,124 SNPs. Quality control of the genotyping data was undertaken using PLINK (55) (version 1.9). No duplicate variants were found. SNPs that were identified as Mendelian inconsistencies (--mendel-multigen) were marked as missing. Gender checks assessed by F-statistic (--check-sex) showed that gender as inferred from 538,771 chromosomal SNPs agreed with reported status. SNPs with low genotyping rates (--geno 0.05), low minor allele frequency (--maf 0.01), and those that failed checks of Hardy-Weinberg Equilibrium (--hwe 1e-8) were excluded. Individuals with low genotype rates (--mind 0.05) and outlying heterozygosity were removed (0.31 - 0.33 included). Relatedness was assessed by high levels of IBD sharing (--genome and --rel-check) and by visualisation of pairs of individuals’ degree of relatedness (through plotting the proportion of loci where the pair shares one allele IBD (Z1) by the proportion of loci where the pair shares zero alleles IBD (Z0)), and two outlier individuals were removed. Ethnicity was assessed via principal components analysis with genotype data from the 1,000 Genome Project (56), which confirmed all participants were of European/CEU origin. Following quality control, 503,221 autosomal SNPs from 1,219 individuals (216 families) were available for SNP-based heritability assessments, of which 999 individuals (196 families; 198 founders and 801 non-founders) had plasma available for lipidomics.

#### 4.3.3 Heritability estimates

SNP-based heritability was estimated using GCTA software (version 1.26.0) (57). A genetic relationship matrix was created from the quality controlled genotyping data and the --reml command was used to estimate variance of the traits explained by the genotyped SNPs. A complementary estimation of pedigree-based heritability was undertaken using the QTDT software (version 2.6.1) (58), by specifying the -we and -veg options to compare an environmental only variance model with a polygenic and environmental variances model. The P-values presented are adjusted for multiple comparisons via Bonferroni correction. The least significant adjusted P-value from the groups of lipid species described are depicted as P_adj_<X.

#### 4.3.4 Genotyping imputation

Following genotyping quality control, 503,221 autosomal SNPs were available to inform imputation. Imputation was performed through the Michigan Imputation Server (version v1.0.4), specifying pre-phasing with Eagle (59) (version 2.3) and imputation by Minimac3 (60) using the European population of the Human Reference Consortium (61) (version hrc.r1.1.2016). Following imputation, duplicate SNPs and SNPs with r^2^<0.8 were removed to generate a final set of 10,652,600 SNPs. Quality control was undertaken for the imputed data on the 999 individuals with lipidomics available, as follows: SNPs that were identified as Mendelian inconsistencies (--mendel-multigen) were marked as missing. SNPs with low call rates (--geno 0.05), low minor allele frequency (--maf 0.05), and those that failed checks of Hardy-Weinberg Equilibrium (--hwe 1e-8) were excluded, resulting in a final count of 5,280,459 SNPs available for genome-wide association analyses.

#### 4.3.5 Family-based genome-wide association studies

Linear mixed modelling approaches were used to account for family structure. Family-based genome-wide association analyses were undertaken for each lipid trait using GCTA software (version 1.26.0), specifying mixed linear model association analyses (--mlma). Genomic control inflation factors from the GWAS analyses can be found in Table S5. The least significant P-values of the significantly associated SNPs (P<5×10^−8^) are depicted as P<X in the manuscript. Significantly associated SNPs were analysed by Ensembl API Client (version 1.1.5 on GRCh37.p13) to identify neighbouring genes. Further analyses were undertaken of the significantly associated SNPs; expression quantitative trait loci (eQTL) were identified using the GTEx eQTL Browser (version 8), assessment of previously identified SNPs from GWAS was undertaken using the GWAS Catalog, review of variants on OMIM, visualisation of variants using UCSC Genome Browser, and assessment of PheWAS with the UK Biobank (62) was undertaken using the Gene Atlas Browser.

#### 4.3.6 Two-sample Mendelian randomisation analysis

Two sample Mendelian randomisation (2SMR) analysis was undertaken in R following the guidelines provided by Davey Smith et al [https://mrcieu.github.io/TwoSampleMR/] (63). Briefly, selected examples of the significant associations identified for each class of lipid were analysed by 2SMR for a number of previously published GWAS of interest. The GWAS significant associations (P<5×10^−8^) identified for NAE species PEA, and ceramide traits CER[N(22)S(19)], CER[N(24)S(16)], and CER[N(24)S(19)]/CER[N(24)DS(19)] ratio, were assessed for coronary artery disease (all), Type-2 Diabetes (CER[N(22)S(19)]), and blood cell counts (CER[N(24)S(16)] and CER[N(24)S(19)]/CER[N(24)DS(19)] ratio). Details on the published GWAS used as outcomes are presented in Table S6. As many GWAS associated SNPs were in linkage disequilibrium, the following SNPs remained in the analysis after the data clumping step; rs324420 (*FAAH*; PEA), rs438568 (*SPTLC3*; CER[N(22)S(19)]), rs7160525 (*SGPP1*; CER[N(24)S(16)]), and rs4653568 (*DEGS1*; CER[N(24)S(19)]/CER[N(24)DS(19)] ratio).

## Acknowledgements and funding support

We are grateful to the families involved in this study. This work was supported by MRC Doctoral Award (MR/K501311/1 to K.M.). K.M. is supported by a University of Manchester President’s Doctoral Scholarship. A.N. is supported in part by the NIHR Manchester Biomedical Research Centre. B.K. is supported by a British Heart Foundation Personal Chair.

## Author contributions

KM: design, analysis and interpretation of data; drafting the article; final approval of the version to be submitted. SW: analysis of data; revising the article; final approval of the version to be submitted. HG: analysis of data; revising the article; final approval of the version to be submitted. HW: acquisition of data; revising the article; final approval of the version to be submitted. MF: analysis and interpretation of data; revising the article; final approval of the version to be submitted. HC: analysis and interpretation of data; revising the article; final approval of the version to be submitted. AN, BK: conception, design, acquisition, and interpretation of data; supervision; revising the article; final approval of the version to be submitted.

## Abbreviations

(NAE): *N*-acyl ethanolamine;
(CER): ceramide;
(eQTL): expression quantitative trait loci;
(h^2^): heritability;
(SNP): single nucleotide polymorphism;
(UPLC-MS/MS): ultra performance liquid chromatography and tandem mass spectrometry;
(AEA): Anandamide (*N*-arachidonoyl ethanolamide);
(DHEA): *N*-docosahexaenoyl ethanolamide;
(DPEA): *N*-docosapentaenoyl ethanolamine;
(LEA): *N*-linoleoyl ethanolamide;
(OEA): *N*-oleoyl ethanolamide;
(PEA): *N*-palmitoyl ethanolamide;
(VEA): *N*-vaccinoyl ethanolamide;
(STEA): *N-*stearoyl ethanolamide;
(HEA): *N*-heptadecanoyl ethanolamide.

---

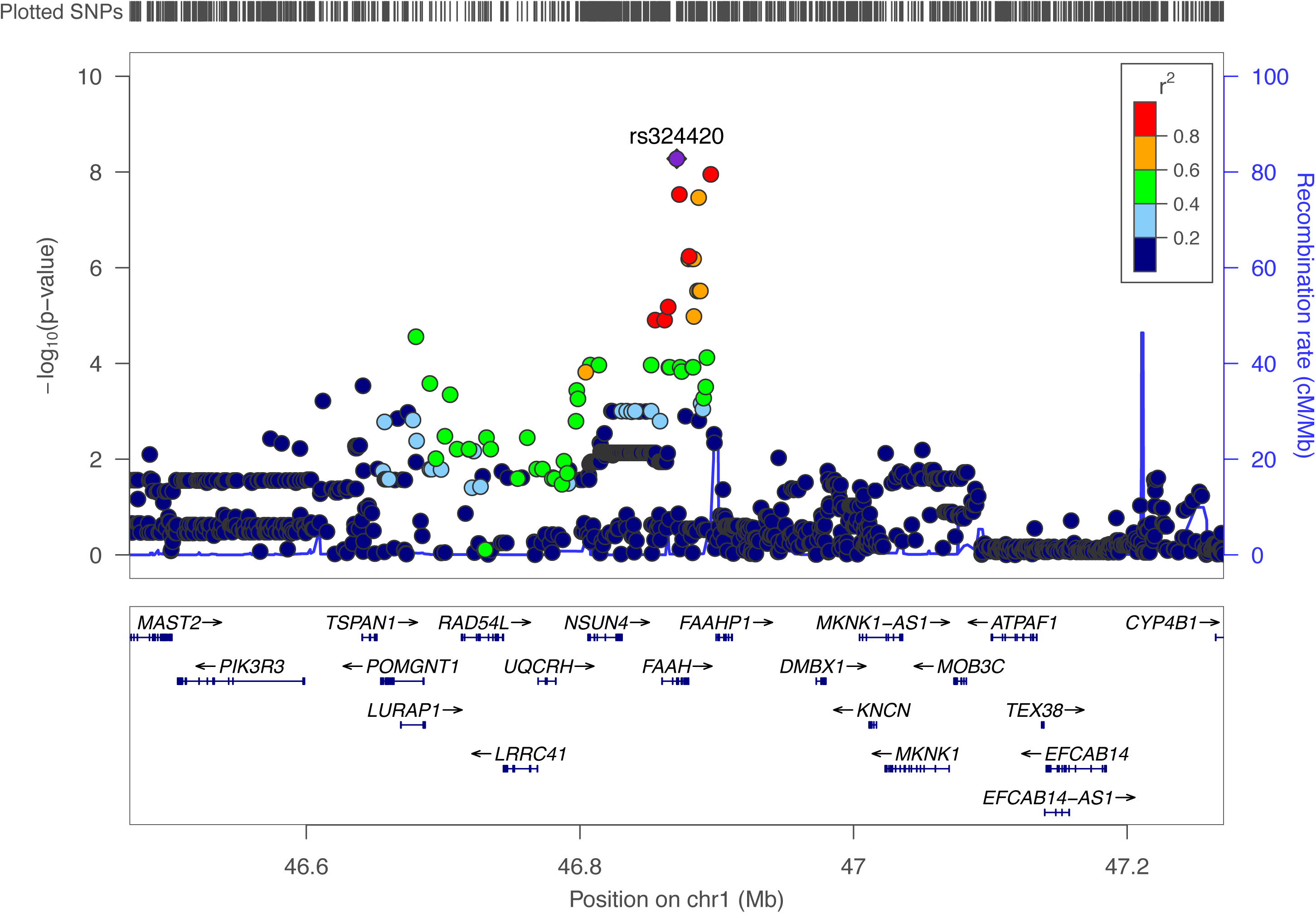

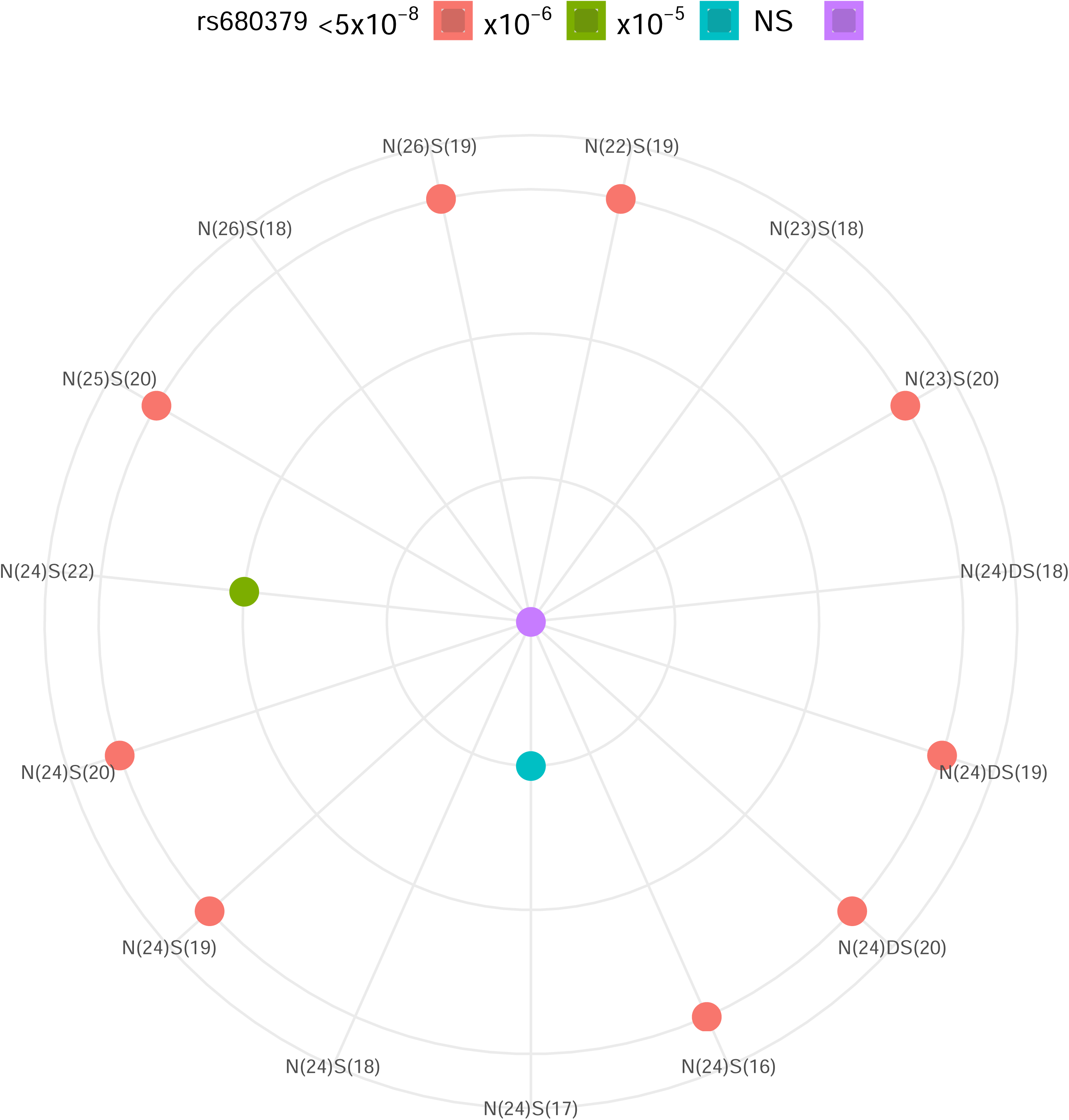

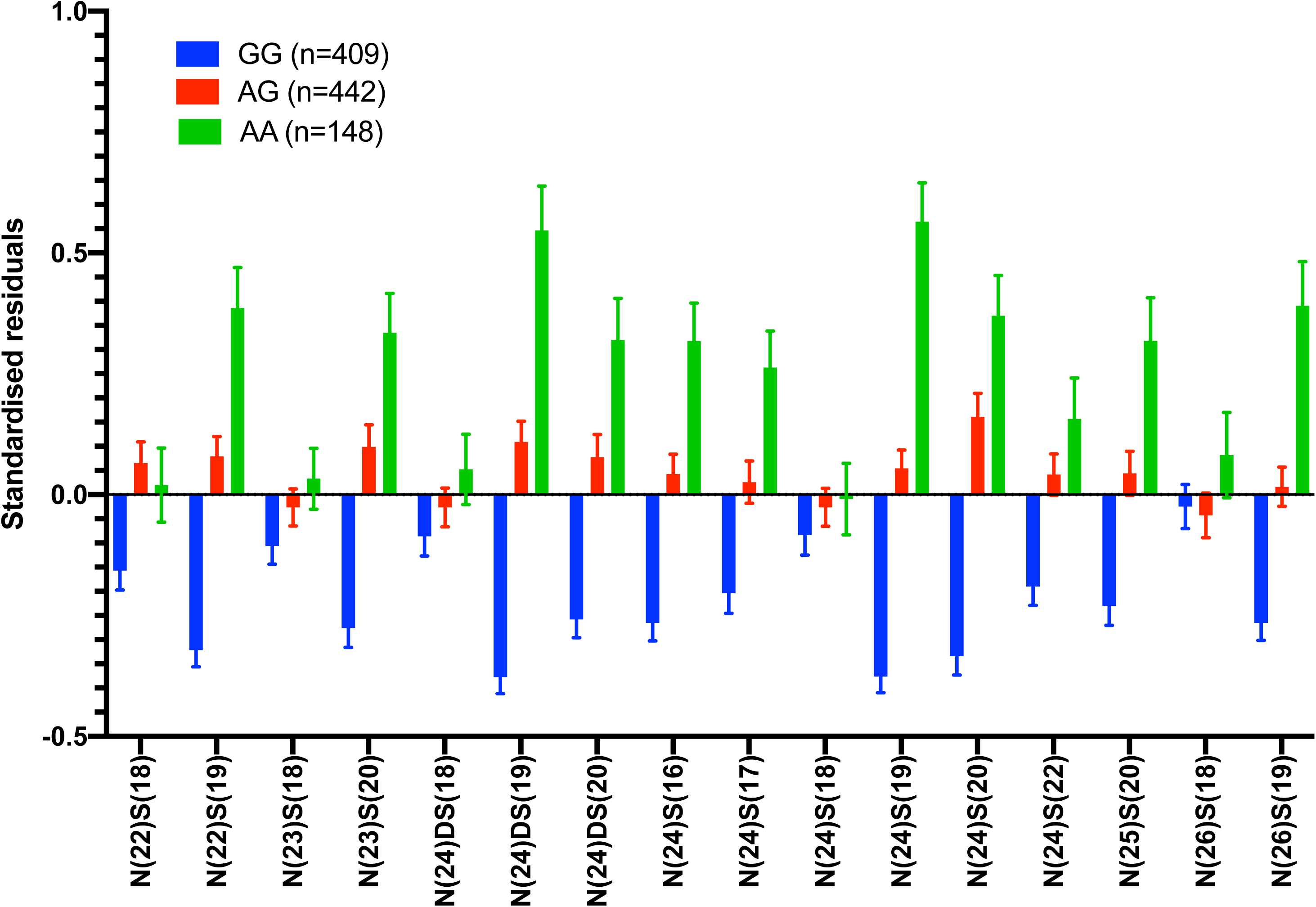

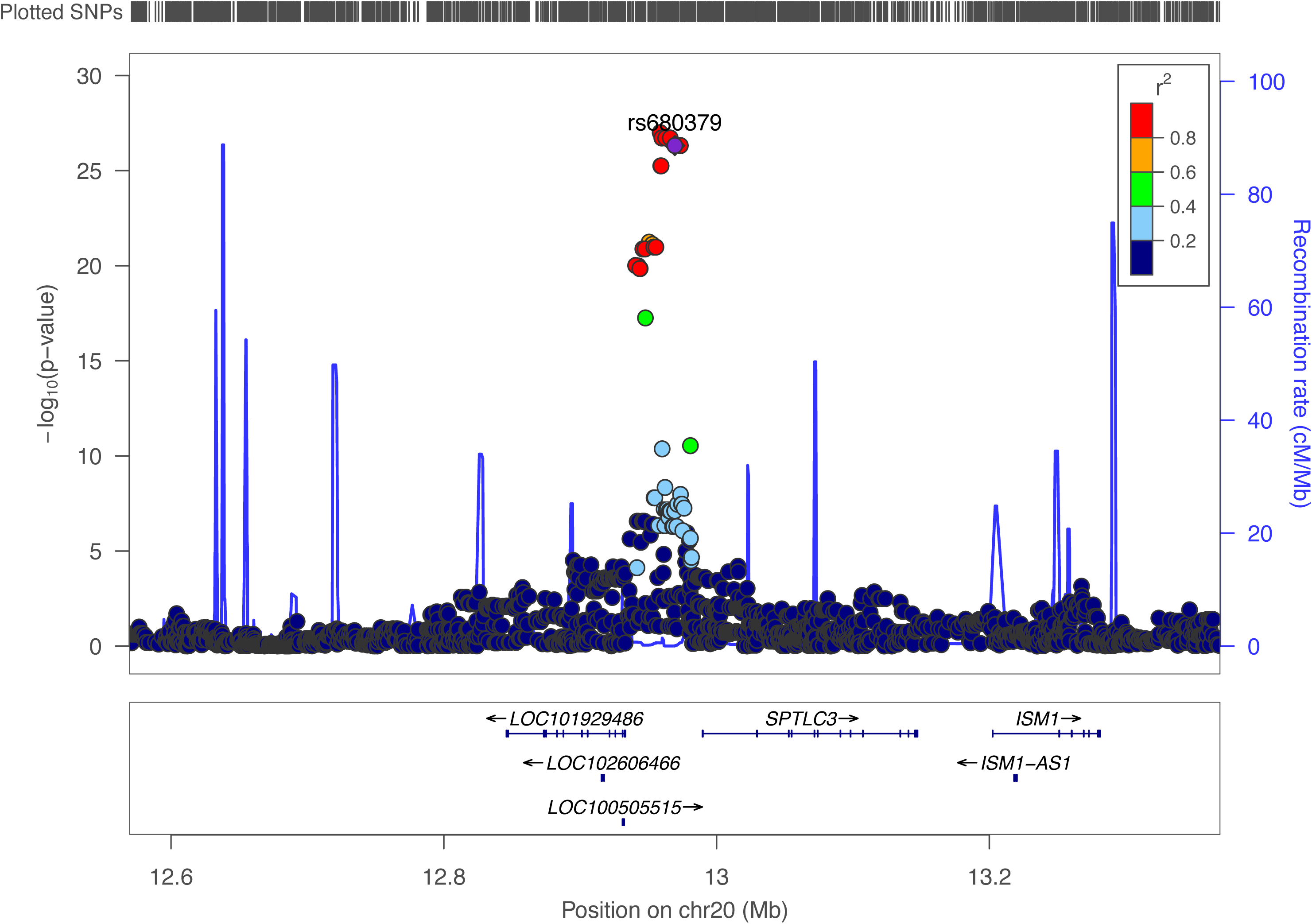

