## Supplementary material for "Heritability and family-based GWAS analyses of the *N*-acyl ethanolamine and ceramide plasma lipidome": Tables and Figures SX

**Supplemental Materials**

**Supplemental methods**

**Quality assessment of plasma NAE species analysis**

Commercially available standards were used for peak referencing of the NAE species. Three pooled plasma samples (quality control; QC) were spiked with commercially available synthetic NAE standards added at a known concentration, and analysed in triplicate with blank ethanol injections run in between samples. Injection variation was analysed by calculation of the mean response and standard deviation of the triplicate injections for each of the three QC samples. The coefficient of variation was calculated, and the mean values are shown in Tables S1. A pooled sample was analysed alongside each batch of clinical samples and a species-specific response value was added to the adjustment model to account for inter-batch variability. An estimation of sample carry-over was calculated by assessing the presence of lipid species in blank injections; there was detectable carryover of NAE species.

**Table S1:** Mean injection coefficient of variation (CV) (n=3 samples) of plasma N-acyl ethanolamine (NAE) species.

| NAE | Mean  Injection  CV (%) |
| --- | --- |
| AEA | 4 |
| PEA | 4 |
| OEA | 6 |
| VEA | 6 |
| LEA | 3 |
| DPEA | 2 |
| DHEA | 6 |
| HEA | 4 |
| STEA | 15 |

**Quality assessment of plasma ceramide species analysis**

Due to lack of commercially available synthetic standards for every ceramide species detected in human plasma, ceramides were analysed via multiple reaction monitoring (MRM) using three transitions per target compound (Table S2). A pooled lipid extract sample was used for the identification of ceramide species, and the coefficient of variation was calculated (Table S3).

**Table S2: Multiple reaction monitoring transitions used for plasma ceramide analysis.** R = fatty acyl chain

| **Ceramide** | ***m/z*** | ***m/z*** | ***m/z*** | ***m/z*** | ***m/z*** |
| --- | --- | --- | --- | --- | --- |
|  | **Precursor** | **Fragment 1** | **Fragment 2** | **Fragment 3** | **Fragment 4** |
|  | **[M+H]^+^** | **[M+H-R-HCHO]^+^** | **[M+H-R-2H_2_O]^+^** | **[M+H-R-H_2_O]^+^** | **[M+H-R]+** |
| CER[N(22)S(18)] | 662.6 | 252.4 | 264.4 | 282.4 |  |
| CER[N(22)S(19)] | 636.6 | 266.4 | 278.4 | 296.4 |  |
| CER[N(23)S(18)] | 636.6 | 252.4 | 264.4 | 282.4 |  |
| CER[N(23)S(20)] | 664.7 | 280.4 | 292.4 | 310.4 |  |
| CER[N(24)DS(18)] | 652.7 |  | 266.4 | 284.4 | 302.4 |
| CER[N(24)DS(19)] | 666.7 |  | 280.4 | 289.4 | 316.4 |
| CER[N(24)DS(20)] | 680.7 |  | 294.4 | 312.4 | 330.4 |
| CER[N(24)S(16)] | 622.6 | 224.4 | 236.4 | 254.4 |  |
| CER[N(24)S(17)] | 636.6 | 238.4 | 250.4 | 268.4 |  |
| CER[N(24)S(18)] | 650.7 | 252.4 | 264.4 | 282.4 |  |
| CER[N(24)S(19)] | 664.7 | 266.4 | 278.4 | 296.4 |  |
| CER[N(24)S(20)] | 678.7 | 280.4 | 292.4 | 310.4 |  |
| CER[N(24)S(22)] | 706.7 | 308.4 | 320.4 | 338.4 |  |
| CER[N(25)S(20)] | 692.7 | 280.4 | 292.4 | 310.4 |  |
| CER[N(26)S(18)] | 678.7 | 252.4 | 264.4 | 282.4 |  |
| CER[N(26)S(19)] | 692.7 | 266.4 | 278.4 | 296.4 |  |

**Table S3: Mean injection coefficient of variation (CV) (n=3 samples) of plasma ceramide species.** *species excluded from the analysis due to high carry-over.

| Ceramide | Mean injection  CV (%) |
| --- | --- |
| CER[N(22)S(18)] | 3 |
| CER[N(22)S(19)] | 11 |
| CER[N(23)S(18)] | 5 |
| CER[N(23)S(20)] | 3 |
| CER[N(24)DS(18)] | 2 |
| CER[N(24)DS(19)] | 6 |
| CER[N(24)DS(20)] | 7 |
| CER[N(24)S(16)] | 5 |
| CER[N(24)S(17)] | 5 |
| CER[N(24)S(18)] | 6 |
| CER[N(24)S(19)] | 7 |
| CER[N(24)S(20)] | 6 |
| CER[N(24)S(22)] | 9 |
| CER[N(25)S(20)] | 7 |
| CER[N(26)S(18)] | 3 |
| CER[N(26)S(19)] | 4 |

**Supplemental Tables**

**Table S4.** Predictors identified from stepwise-multiple linear regression.

qc, quality control sample; Batch, mass spectrometry batch; Age, age at enrolment; Abnormality, presence of white blood cells, red blood cells or other sample abnormality; sumorratio, summation or ratio of multiple species described in Table S4.

| **Class** | **Lipid** | **Predictors** |
| --- | --- | --- |
| CER | N(22)S(18) | N22_S18_qc + Batch + Abnormality + Age + Sex + Hypertension + Age2 |
| CER | N(22)S(19) | N22_S19_qc + Batch + Abnormality + Cholesterol |
| CER | N(23)S(18) | N23_S18_qc + Batch + Age2 + Age |
| CER | N(23)S(20) | N23_S20_qc + Batch + Sex + Age2 |
| CER | N(24)DS(18) | N24_DS18_qc + Batch + Sex + Age + BMI + Cholesterol |
| CER | N(24)DS(19) | N24_DS19_qc + Batch + Sex + BMI + Cholesterol |
| CER | N(24)DS(20) | N24_DS20_qc + Batch + Abnormality + Sex + Hypertension + Age + Age2 + BMI + Cholesterol |
| CER | N(24)S(16) | N24_S16_qc + Batch + Abnormality + Sex + Age + Age2 |
| CER | N(24)S(17) | N24_S17_qc + Batch + Sex + Age + Age2 |
| CER | N(24)S(18) | N24_S18_qc + Batch + Age + Age2 + Cholesterol |
| CER | N(24)S(19) | N24_S19_qc + Batch + Abnormality |
| CER | N(24)S(20) | N24_S20_qc + Batch + Abnormality + Sex + Hypertension + Age + Age2 + BMI + Cholesterol |
| CER | N(24)S(22) | N24_S22_qc + Sex + Age2 + BMI |
| CER | N(25)S(20) | N25_S20_qc + Age + BMI + Cholesterol |
| CER | N(26)S(18) | N26_S18_qc + Batch + Sex + Age + Age2 + Cholesterol |
| CER | N(26)S(19) | N26_S19_qc + Sex + Age2 + BMI + Cholesterol |
| NAE | AEA | AEA_qc + Cholesterol + BMI + Batch |
| NAE | DHEA | DHEA_qc + Cholesterol + Age2 + Batch |
| NAE | DPEA | DPEA_qc + Cholesterol + BMI + Batch + Sex + Hypertension + Age2 + Age + Abnormality |
| NAE | OEA | OEA_qc + Cholesterol + Batch + BMI |
| NAE | LEA | LEA_qc + Cholesterol + Batch + Age2 |
| NAE | PEA | PEA_qc + Cholesterol |
| NAE | VEA | VEA_qc + Cholesterol + Batch + Sex + Hypertension |
| NAE | STEA | STEA_qc + Cholesterol + Batch + Sex + BMI |
| NAE | HEA | HEA_qc + Cholesterol + Batch |

**Table S5.** Genomic inflation factors (GIF) from GWAS results per trait.

| **Class** | **Lipid** | **GIF** |
| --- | --- | --- |
| CER | N(22)S(18) | 1.0042 |
| CER | N(22)S(19) | 0.9893 |
| CER | N(23)S(18) | 1.0000 |
| CER | N(23)S(20) | 1.0046 |
| CER | N(24)DS(18) | 1.0044 |
| CER | N(24)DS(19) | 0.9952 |
| CER | N(24)DS(20) | 1.0045 |
| CER | N(24)S(16) | 1.0012 |
| CER | N(24)S(17) | 1.0037 |
| CER | N(24)S(18) | 1.0036 |
| CER | N(24)S(19) | 0.9944 |
| CER | N(24)S(20) | 0.9907 |
| CER | N(24)S(22) | 1.0036 |
| CER | N(25)S(20) | 1.0061 |
| CER | N(26)S(18) | 1.0017 |
| CER | N(26)S(19) | 0.9939 |
| CER | n24s19ratio | 0.9937 |
| CER | s19_sum | 0.9954 |
| CER | s20_sum | 0.9945 |
| NAE | AEA | 0.9984 |
| NAE | DHEA | 0.9916 |
| NAE | DPEA | 1.0062 |
| NAE | LEA | 0.9910 |
| NAE | OEA | 0.9916 |
| NAE | PEA | 0.9904 |
| NAE | VEA | 0.9921 |
| NAE | STEA | 0.9984 |
| NAE | HEA | 0.9953 |
| NAE | sumEA | 0.9961 |

**Table S6.** List of GWAS assessed by 2SMR analysis for relationship with the SNPs identified at GWAS to associate with the lipid species.

id, 2SMR software ID; pmid, PubMed ID; pop, population type.

| **author** | **consortium** | **Filename** | **id** | **ncase** | **ncontrol** | **nsnp** | **pmid** | **pop** | **sample**  **size** | **sex** | **trait** | **unit** | **year** |
| --- | --- | --- | --- | --- | --- | --- | --- | --- | --- | --- | --- | --- | --- |
| Nikpay | CARDIoGRAMplusC4D | cad.add.160614.website.txt.tab | 7 | 60801 | 123504 | 9455779 | 26343387 | Mixed | 184305 | Males and females | Coronary heart disease | log odds | 2015 |
| Mahajan A | DIAGRAM | diagram.mega-meta.txt.tab.all_pos | 23 | 26488 | 83964 | 2915012 | 24509480 | Mixed | 110452 | Males and females | Type 2 diabetes | log odds | 2014 |
| van der Harst P | HaemGen | HaemGenRBC_RBC.txt.tab | 275 | NA | NA | 2589455 | 23222517 | Mixed | 66214 | Males and females | Red blood cell count | 10^12/L | 2012 |
| Gieger C | HaemGen | PLT.txt.tab | 1008 | NA | NA | 2703394 | 22139419 | European | 66867 | Males and females | Platelet count | 1x10^9/l | 2011 |
| Astle W | UK Biobank+INTERVAL+UK BiLEVE | 27863252-GCST004599-EFO_0004584-Build37.tab.gz | 1247 | NA | NA | 29462512 | 27863252 | European | 164454 | Males and females | Mean platelet volume | SD | 2016 |
| Astle W | UK Biobank+INTERVAL+UK BiLEVE | 27863252-GCST004600-EFO_0007991-Build37.tab.gz | 1248 | NA | NA | 29483746 | 27863252 | European | 172378 | Males and females | Eosinophil percentage of white cells | SD | 2016 |
| Astle W | UK Biobank+INTERVAL+UK BiLEVE | 27863252-GCST004601-EFO_0004305-Build37.tab.gz | 1249 | NA | NA | 29486177 | 27863252 | European | 172952 | Males and females | Red blood cell count | SD | 2016 |
| Astle W | UK Biobank+INTERVAL+UK BiLEVE | 27863252-GCST004602-EFO_0004526-Build37.tab.gz | 1250 | NA | NA | 29483230 | 27863252 | European | 172433 | Males and females | Mean corpuscular volume | SD | 2016 |
| Astle W | UK Biobank+INTERVAL+UK BiLEVE | 27863252-GCST004603-EFO_0004309-Build37.tab.gz | 1251 | NA | NA | 29465077 | 27863252 | European | 166066 | Males and females | Platelet count | SD | 2016 |
| Astle W | UK Biobank+INTERVAL+UK BiLEVE | 27863252-GCST004604-EFO_0004348-Build37.tab.gz | 1252 | NA | NA | 29484426 | 27863252 | European | 173039 | Males and females | Hematocrit | SD | 2016 |
| Astle W | UK Biobank+INTERVAL+UK BiLEVE | 27863252-GCST004605-EFO_0004528-Build37.tab.gz | 1253 | NA | NA | 29486070 | 27863252 | European | 172851 | Males and females | Mean corpuscular hemoglobin concentration | SD | 2016 |
| Astle W | UK Biobank+INTERVAL+UK BiLEVE | 27863252-GCST004606-EFO_0004842-Build37.tab.gz | 1254 | NA | NA | 29485759 | 27863252 | European | 172275 | Males and females | Eosinophil counts | SD | 2016 |
| Astle W | UK Biobank+INTERVAL+UK BiLEVE | 27863252-GCST004607-EFO_0007985-Build37.tab.gz | 1255 | NA | NA | 29463315 | 27863252 | European | 164339 | Males and females | Plateletcrit | SD | 2016 |
| Astle W | UK Biobank+INTERVAL+UK BiLEVE | 27863252-GCST004608-EFO_0007997-Build37.tab.gz | 1256 | NA | NA | 29480509 | 27863252 | European | 169545 | Males and females | Granulocyte percentage of myeloid white cells | SD | 2016 |
| Astle W | UK Biobank+INTERVAL+UK BiLEVE | 27863252-GCST004609-EFO_0007989-Build37.tab.gz | 1257 | NA | NA | 29481677 | 27863252 | European | 170494 | Males and females | Monocyte percentage of white cells | SD | 2016 |
| Astle W | UK Biobank+INTERVAL+UK BiLEVE | 27863252-GCST004610-EFO_0004308-Build37.tab.gz | 1258 | NA | NA | 29485724 | 27863252 | European | 172435 | Males and females | White blood cell count | SD | 2016 |
| Astle W | UK Biobank+INTERVAL+UK BiLEVE | 27863252-GCST004611-EFO_0007986-Build37.tab.gz | 1259 | NA | NA | 29480430 | 27863252 | European | 170761 | Males and females | High light scatter reticulocyte count | SD | 2016 |
| Astle W | UK Biobank+INTERVAL+UK BiLEVE | 27863252-GCST004612-EFO_0007986-Build37.tab.gz | 1260 | NA | NA | 29480170 | 27863252 | European | 170763 | Males and females | High light scatter reticulocyte percentage of red cells | SD | 2016 |
| Astle W | UK Biobank+INTERVAL+UK BiLEVE | 27863252-GCST004613-EFO_0004833-Build37.tab.gz | 1261 | NA | NA | 29482518 | 27863252 | European | 170384 | Males and females | Sum neutrophil eosinophil counts | SD | 2016 |
| Astle W | UK Biobank+INTERVAL+UK BiLEVE | 27863252-GCST004614-EFO_0007987-Build37.tab.gz | 1262 | NA | NA | 29481601 | 27863252 | European | 169822 | Males and females | Granulocyte count | SD | 2016 |
| Astle W | UK Biobank+INTERVAL+UK BiLEVE | 27863252-GCST004615-EFO_0004509-Build37.tab.gz | 1263 | NA | NA | 29483564 | 27863252 | European | 172925 | Males and females | Hemoglobin concentration | SD | 2016 |
| Astle W | UK Biobank+INTERVAL+UK BiLEVE | 27863252-GCST004616-EFO_0007984-Build37.tab.gz | 1264 | NA | NA | 29461105 | 27863252 | European | 164433 | Males and females | Platelet distribution width | SD | 2016 |
| Astle W | UK Biobank+INTERVAL+UK BiLEVE | 27863252-GCST004617-EFO_0007996-Build37.tab.gz | 1265 | NA | NA | 29482376 | 27863252 | European | 170536 | Males and females | Eosinophil percentage of granulocytes | SD | 2016 |
| Astle W | UK Biobank+INTERVAL+UK BiLEVE | 27863252-GCST004618-EFO_0005090-Build37.tab.gz | 1266 | NA | NA | 29484325 | 27863252 | European | 171846 | Males and females | White blood cell count (basophil) | SD | 2016 |
| Astle W | UK Biobank+INTERVAL+UK BiLEVE | 27863252-GCST004619-EFO_0007986-Build37.tab.gz | 1267 | NA | NA | 29480520 | 27863252 | European | 170690 | Males and females | Reticulocyte fraction of red cells | SD | 2016 |
| Astle W | UK Biobank+INTERVAL+UK BiLEVE | 27863252-GCST004620-EFO_0004833-Build37.tab.gz | 1268 | NA | NA | 29480759 | 27863252 | European | 170143 | Males and females | Sum basophil neutrophil counts | SD | 2016 |
| Astle W | UK Biobank+INTERVAL+UK BiLEVE | 27863252-GCST004621-EFO_0005192-Build37.tab.gz | 1269 | NA | NA | 29484006 | 27863252 | European | 171529 | Males and females | Red cell distribution width | SD | 2016 |
| Astle W | UK Biobank+INTERVAL+UK BiLEVE | 27863252-GCST004622-EFO_0007986-Build37.tab.gz | 1270 | NA | NA | 29479992 | 27863252 | European | 170641 | Males and females | Reticulocyte count | SD | 2016 |
| Astle W | UK Biobank+INTERVAL+UK BiLEVE | 27863252-GCST004623-EFO_0007994-Build37.tab.gz | 1271 | NA | NA | 29482650 | 27863252 | European | 170672 | Males and females | Neutrophil percentage of granulocytes | SD | 2016 |
| Astle W | UK Biobank+INTERVAL+UK BiLEVE | 27863252-GCST004624-EFO_0005090-Build37.tab.gz | 1272 | NA | NA | 29485063 | 27863252 | European | 171771 | Males and females | Sum eosinophil basophil counts | SD | 2016 |
| Astle W | UK Biobank+INTERVAL+UK BiLEVE | 27863252-GCST004625-EFO_0005091-Build37.tab.gz | 1273 | NA | NA | 29482454 | 27863252 | European | 170721 | Males and females | Monocyte count | SD | 2016 |
| Astle W | UK Biobank+INTERVAL+UK BiLEVE | 27863252-GCST004626-EFO_0007988-Build37.tab.gz | 1274 | NA | NA | 29478559 | 27863252 | European | 169219 | Males and females | Myeloid white cell count | SD | 2016 |
| Astle W | UK Biobank+INTERVAL+UK BiLEVE | 27863252-GCST004627-EFO_0004587-Build37.tab.gz | 1275 | NA | NA | 29484106 | 27863252 | European | 171643 | Males and females | Lymphocyte counts | SD | 2016 |
| Astle W | UK Biobank+INTERVAL+UK BiLEVE | 27863252-GCST004628-EFO_0007986-Build37.tab.gz | 1276 | NA | NA | 29479929 | 27863252 | European | 170548 | Males and females | Immature fraction of reticulocytes | SD | 2016 |
| Astle W | UK Biobank+INTERVAL+UK BiLEVE | 27863252-GCST004629-EFO_0004833-Build37.tab.gz | 1277 | NA | NA | 29481373 | 27863252 | European | 170702 | Males and females | Neutrophil count | SD | 2016 |

**Table S7:** Lipid species, corresponding class, summary statistics and description of the species.

The concentrations are provided before adjustment or outlier removal for a total of 999 samples analysed from 196 families.

| **Lipid** | **Class** | **unit** | **N** | **Mean** | **SD** | **Description of lipid species** |
| --- | --- | --- | --- | --- | --- | --- |
| N(22)S(18) | CER | pmol/ml | 999 | 128 | 74 | Non-hydroxy fatty acid and sphingosine bas |
| N(22)S(19) | CER | pmol/ml | 999 | 32 | 22 | Non-hydroxy fatty acid and sphingosine base |
| N(23)S(18) | CER | pmol/ml | 999 | 866 | 365 | Non-hydroxy fatty acid and sphingosine base |
| N(23)S(20) | CER | pmol/ml | 999 | 51 | 16 | Non-hydroxy fatty acid and sphingosine base |
| N(24)DS(18) | CER | pmol/ml | 999 | 175 | 112 | Non-hydroxy fatty acid and dihydrosphingosine base |
| N(24)DS(19) | CER | pmol/ml | 999 | 66 | 42 | Non-hydroxy fatty acid and dihydrosphingosine base |
| N(24)DS(20) | CER | pmol/ml | 998 | 40 | 22 | Non-hydroxy fatty acid and dihydrosphingosine base |
| N(24)S(16) | CER | pmol/ml | 999 | 49 | 30 | Non-hydroxy fatty acid and sphingosine base |
| N(24)S(17) | CER | pmol/ml | 999 | 236 | 106 | Non-hydroxy fatty acid and sphingosine base |
| N(24)S(18) | CER | pmol/ml | 999 | 2724 | 1290 | Non-hydroxy fatty acid and sphingosine base |
| N(24)S(19) | CER | pmol/ml | 999 | 1068 | 486 | Non-hydroxy fatty acid and sphingosine base |
| N(24)S(20) | CER | pmol/ml | 999 | 249 | 93 | Non-hydroxy fatty acid and sphingosine base |
| N(24)S(22) | CER | pmol/ml | 999 | 45 | 31 | Non-hydroxy fatty acid and sphingosine base |
| N(25)S(20) | CER | pmol/ml | 999 | 38 | 19 | Non-hydroxy fatty acid and sphingosine base |
| N(26)S(18) | CER | pmol/ml | 999 | 706 | 219 | Non-hydroxy fatty acid and sphingosine base |
| N(26)S(19) | CER | pmol/ml | 999 | 108 | 74 | Non-hydroxy fatty acid and sphingosine base |
| AEA | NAE | pg/ml | 998 | 352 | 334 | Anandamide (N-arachidonoyl ethanolamide) |
| DHEA | NAE | pg/ml | 996 | 349 | 290 | N-docosahexaenoyl ethanolamide |
| DPEA | NAE | pg/ml | 990 | 22 | 17 | N-docosapentaenoyl ethanolamine |
| LEA | NAE | pg/ml | 999 | 619 | 511 | N-linoleoyl ethanolamide |
| OEA | NAE | pg/ml | 999 | 568 | 531 | N-oleoyl ethanolamide |
| PEA | NAE | pg/ml | 999 | 1884 | 1356 | N-palmitoyl ethanolamide |
| VEA | NAE | pg/ml | 999 | 252 | 259 | N-vaccinoyl ethanolamide |
| HEA | NAE | pg/ml | 968 | 24 | 19 | N-heptadecanoyl ethanolamide |
| STEA | NAE | pg/ml | 999 | 495 | 447 | N-stearoyl ethanolamide |

**Table S8.** Heritability of plasma lipid species measured.

Heritability was estimated using GCTA SNP-based software using the genotyping data and QTDT pedigree-based software. h^2^, estimated heritability; SE, standard error; ChiSq, chi-squared; n, number of individuals included in analysis after outlier removal; P-adj, P-value after Bonferroni adjustment of the P-value for multiple testing (12 for CER measured, 7 for NAE measures). A GCTA P-value of 0 is represented as a P-value result of <6.11x10^-16^.

|  |  | **QTDT** | | | | | **GCTA** | | | | |
| --- | --- | --- | --- | --- | --- | --- | --- | --- | --- | --- | --- |
| **Class** | **Lipid** | **h^2^** | **ChiSq** | **n** | **P-value** | **P-adj** | **h^2^** | **SE** | **n** | **P-value** | **P-adj** |
| CER | N(22)S(18) | 0.49 | 98 | 993 | 5.00E-23 | 1.50E-21 | 0.48 | 0.059 | 993 | 6.11E-16 | 1.83E-14 |
| CER | N(22)S(19) | 0.38 | 64 | 992 | 1.00E-15 | 1.10E-14 | 0.36 | 0.057 | 992 | 6.11E-15 | 6.72E-14 |
| CER | N(23)S(18) | 0.40 | 68 | 992 | 2.00E-16 | 6.00E-15 | 0.39 | 0.057 | 992 | 1.11E-16 | 3.33E-15 |
| CER | N(23)S(20) | 0.53 | 91 | 994 | 1.00E-21 | 1.10E-20 | 0.54 | 0.061 | 994 | 6.11E-16 | 6.72E-15 |
| CER | N(24)DS(18) | 0.40 | 69 | 992 | 8.00E-17 | 2.40E-15 | 0.39 | 0.060 | 992 | 5.55E-17 | 1.67E-15 |
| CER | N(24)DS(19) | 0.45 | 73 | 994 | 1.00E-17 | 1.10E-16 | 0.46 | 0.062 | 994 | 6.11E-16 | 6.72E-15 |
| CER | N(24)DS(20) | 0.52 | 107 | 993 | 6.00E-25 | 6.60E-24 | 0.52 | 0.059 | 993 | 6.11E-16 | 6.72E-15 |
| CER | N(24)S(16) | 0.57 | 150 | 992 | 2.00E-34 | 2.20E-33 | 0.57 | 0.055 | 992 | 6.11E-16 | 6.72E-15 |
| CER | N(24)S(17) | 0.42 | 80 | 994 | 3.00E-19 | 3.30E-18 | 0.44 | 0.057 | 994 | 6.11E-16 | 6.72E-15 |
| CER | N(24)S(18) | 0.47 | 101 | 991 | 1.00E-23 | 1.10E-22 | 0.46 | 0.056 | 991 | 6.11E-16 | 6.72E-15 |
| CER | N(24)S(19) | 0.46 | 72 | 991 | 2.00E-17 | 2.20E-16 | 0.38 | 0.061 | 993 | 9.94E-15 | 1.09E-13 |
| CER | N(24)S(20) | 0.54 | 102 | 996 | 5.00E-24 | 5.50E-23 | 0.55 | 0.059 | 996 | 6.11E-16 | 6.72E-15 |
| CER | N(24)S(22) | 0.55 | 93 | 992 | 6.00E-22 | 6.60E-21 | 0.55 | 0.061 | 992 | 6.11E-16 | 6.72E-15 |
| CER | N(25)S(20) | 0.62 | 126 | 994 | 3.00E-29 | 3.30E-28 | 0.6 | 0.059 | 994 | 6.11E-16 | 6.72E-15 |
| CER | N(26)S(18) | 0.54 | 115 | 995 | 7.00E-27 | 2.10E-25 | 0.54 | 0.059 | 995 | 6.11E-16 | 1.83E-14 |
| CER | N(26)S(19) | 0.43 | 57 | 991 | 4.00E-14 | 4.40E-13 | 0.42 | 0.063 | 991 | 3.24E-14 | 3.56E-13 |
| NAE | AEA | 0.48 | 87 | 994 | 1.00E-20 | 7.00E-20 | 0.46 | 0.060 | 994 | 6.11E-16 | 4.28E-15 |
| NAE | DHEA | 0.56 | 132 | 990 | 2.00E-30 | 1.40E-29 | 0.54 | 0.057 | 990 | 6.11E-16 | 4.28E-15 |
| NAE | DPEA | 0.54 | 99 | 986 | 2.00E-23 | 1.40E-22 | 0.52 | 0.060 | 986 | 6.11E-16 | 4.28E-15 |
| NAE | LEA | 0.46 | 117 | 994 | 3.00E-27 | 2.10E-26 | 0.45 | 0.053 | 994 | 6.11E-16 | 4.28E-15 |
| NAE | OEA | 0.45 | 84 | 994 | 4.00E-20 | 2.80E-19 | 0.45 | 0.058 | 994 | 6.11E-16 | 4.28E-15 |
| NAE | PEA | 0.54 | 140 | 993 | 2.00E-32 | 1.40E-31 | 0.53 | 0.054 | 993 | 6.11E-16 | 4.28E-15 |
| NAE | VEA | 0.49 | 105 | 994 | 1.00E-24 | 7.00E-24 | 0.5 | 0.058 | 994 | 6.11E-16 | 4.28E-15 |
| NAE | STEA | 0.62 | 209 | 994 | 2.00E-47 | 2.20E-46 | 0.60 | 0.049 | 994 | 6.11E-16 | 6.72E-15 |
| NAE | HEA | 0.69 | 232 | 964 | 2.00E-52 | 2.20E-51 | 0.68 | 0.050 | 964 | 6.11E-16 | 6.72E-15 |

**Table S9.** Significant GWAS associations for *N*-acyl ethanolamine species.

A) Description of the GWAS significant associations identified. B) Description of the SNPs using the Ensembl API Client. C) Summary of the information on eQTL status as identified using the GTEX browser, including a section specifying whole blood only, GWAS Catalog information, and Gene Atlas PheWAS.

| **A** |  |  |  |  |  |  |  |  |  |
| --- | --- | --- | --- | --- | --- | --- | --- | --- | --- |
| **Lipid** | **Chr** | **SNP** | **Position** | **A1** | **A2** | **MAF** | **Beta** | **SE** | **P-value** |
| DHEA | 1 | rs324420 | 46870761 | A | C | 0.20 | 0.30 | 0.053 | 2.15E-08 |
| DHEA | 1 | rs324422 | 46886782 | T | C | 0.23 | 0.28 | 0.050 | 2.76E-08 |
| DHEA | 1 | rs324418 | 46872698 | G | A | 0.22 | 0.28 | 0.052 | 4.36E-08 |
| LEA | 1 | rs324420 | 46870761 | A | C | 0.20 | 0.31 | 0.055 | 1.01E-08 |
| LEA | 1 | rs1571138 | 46895641 | A | G | 0.20 | 0.31 | 0.055 | 2.24E-08 |
| LEA | 1 | rs324422 | 46886782 | T | C | 0.23 | 0.29 | 0.052 | 2.32E-08 |
| PEA | 1 | rs324420 | 46870761 | A | C | 0.21 | 0.30 | 0.051 | 5.30E-09 |
| PEA | 1 | rs1571138 | 46895641 | A | G | 0.20 | 0.29 | 0.051 | 1.13E-08 |
| PEA | 1 | rs324418 | 46872698 | G | A | 0.22 | 0.27 | 0.050 | 2.96E-08 |
| PEA | 1 | rs324422 | 46886782 | T | C | 0.23 | 0.27 | 0.048 | 3.44E-08 |
| VEA | 1 | rs324420 | 46870761 | A | C | 0.20 | 0.31 | 0.049 | 1.24E-10 |
| VEA | 1 | rs1571138 | 46895641 | A | G | 0.20 | 0.32 | 0.049 | 1.25E-10 |
| VEA | 1 | rs324418 | 46872698 | G | A | 0.22 | 0.29 | 0.048 | 1.15E-09 |
| VEA | 1 | rs11584511 | 46892811 | T | C | 0.14 | 0.34 | 0.058 | 3.35E-09 |
| VEA | 1 | rs324422 | 46886782 | T | C | 0.23 | 0.27 | 0.047 | 4.66E-09 |
| VEA | 1 | rs10489770 | 46807597 | A | G | 0.14 | 0.32 | 0.058 | 4.22E-08 |
| VEA | 1 | rs72677586 | 46813848 | A | G | 0.14 | 0.32 | 0.058 | 4.22E-08 |
| VEA | 1 | rs10890392 | 46852061 | G | A | 0.14 | 0.32 | 0.058 | 4.22E-08 |
| sumEA | 1 | rs324420 | 46870761 | A | C | 0.21 | 0.32 | 0.053 | 1.36E-09 |
| sumEA | 1 | rs1571138 | 46895641 | A | G | 0.20 | 0.31 | 0.053 | 2.62E-09 |
| sumEA | 1 | rs324418 | 46872698 | G | A | 0.22 | 0.29 | 0.051 | 1.84E-08 |
| sumEA | 1 | rs324422 | 46886782 | T | C | 0.23 | 0.28 | 0.050 | 2.10E-08 |

| **B** |  |  |  |  |  |  |  |
| --- | --- | --- | --- | --- | --- | --- | --- |
| **SNPID** | **Associated**  **Gene ID** | **Associated**  **Transcript ID** | **Associated**  **Gene Name** | **Associated**  **Gene Type** | **Impact**  **Rating** | **Variant**  **Allele** | **Consequence**  **Terms** |
| rs324420 | ENSG00000117480 | ENST00000243167 | FAAH | Protein coding | MODERATE | A | Missense variant |
| rs324422 | (Intergenic) | (Intergenic) | (Intergenic) | (Intergenic) | MODIFIER | T | Intergenic variant |
| rs324418 | ENSG00000117480 | ENST00000243167 | FAAH | Protein coding | MODIFIER | G | Intron variant |
| rs1571138 | ENSG00000232022 | ENST00000446499 | FAAHP1 | pseudogene | MODIFIER | G | upstream |
| rs11584511 | ENSG00000232022 | ENST00000446499 | FAAHP1 | pseudogene | MODIFIER | T | upstream |
| rs10489770 | ENSG00000117481 | ENST00000307089 | NSUN4 | NMD | MODIFIER | A | intron_variant |
| rs72677586 | ENSG00000117481 | ENST00000307089 | NSUN4 | NMD | MODIFIER | A | intron_variant |
| rs10890392 | (Intergenic) | (Intergenic) | (Intergenic) | (Intergenic) | MODIFIER | G | intergenic_variant |

| **C** |  |  |  |  |  |  |
| --- | --- | --- | --- | --- | --- | --- |
| **SNPID** | **eQTL** | **Tissue** | **Other** | **GTEx whole blood** | **GWAS Catalog** | **Gene Atlas** |
| rs324420 | FAAH | Multiple | FAAHP1, LURAP1, NSUN4, RAD54L, MKNK1 | FAAH, NSUN4 | X | X |
| rs324422 | FAAH | Multiple | FAAHP1, LURAP1, NSUN4, RAD54L, MKNK1, MOB3C | FAAH, NSUN4, MOB3C | X | X |
| rs324418 | FAAH | Multiple | FAAHP1, LURAP1, NSUN4, RAD54L, MKNK1 | FAAH, NSUN4 | X | X |
| rs1571138 | FAAH | Multiple | FAAHP1, LURAP1, NSUN4, RAD54L, MKNK1 | FAAH, NSUN4 | X | X |
| rs11584511 | FAAH | Multiple | FAAHP1, LURAP1, NSUN4, RAD54L, MKNK1, UQCRH | FAAH, NSUN4 | X | X |
| rs10489770 | FAAH | Multiple | FAAHP1, LURAP1, NSUN4, RAD54L, MKNK1, UQCRH | FAAH | X | X |
| rs72677586 | FAAH | Multiple | FAAHP1, LURAP1, NSUN4, RAD54L, MKNK1, UQCRH | FAAH | X | X |
| rs10890392 | FAAH | Multiple | FAAHP1, LURAP1, NSUN4, RAD54L, MKNK1, UQCRH | FAAH | X | X |

**Table S10.** Significant GWAS associations for ceramides and related sphingolipid species.

Description of the GWAS significant associations identified.

| **Lipid** | **Chr** | **SNP** | **Position** | **A1** | **A2** | **MAF** | **Beta** | **SE** | **P-value** |
| --- | --- | --- | --- | --- | --- | --- | --- | --- | --- |
| N22S19 | 20 | rs438568 | 12958687 | A | G | 0.36 | 0.37 | 0.044 | 7.59E-18 |
| N22S19 | 20 | rs1321940 | 12959885 | A | G | 0.37 | 0.37 | 0.044 | 1.43E-17 |
| N22S19 | 20 | rs364585 | 12962718 | A | G | 0.37 | 0.37 | 0.044 | 1.43E-17 |
| N22S19 | 20 | rs168622 | 12966089 | T | G | 0.37 | 0.37 | 0.044 | 1.43E-17 |
| N22S19 | 20 | rs680379 | 12969400 | A | G | 0.37 | 0.37 | 0.043 | 2.91E-17 |
| N22S19 | 20 | rs686548 | 12973521 | A | T | 0.37 | 0.37 | 0.043 | 2.91E-17 |
| N22S19 | 20 | rs4814175 | 12959094 | A | T | 0.37 | 0.36 | 0.044 | 1.40E-16 |
| N22S19 | 20 | rs4814176 | 12959398 | T | C | 0.37 | 0.36 | 0.044 | 1.40E-16 |
| N22S19 | 20 | rs2327452 | 12952964 | A | C | 0.32 | 0.34 | 0.046 | 1.30E-13 |
| N22S19 | 20 | rs3848746 | 12950606 | A | G | 0.32 | 0.34 | 0.046 | 1.48E-13 |
| N22S19 | 20 | rs2327451 | 12953934 | C | A | 0.32 | 0.34 | 0.046 | 1.61E-13 |
| N22S19 | 20 | rs4508668 | 12955601 | T | C | 0.32 | 0.34 | 0.046 | 1.61E-13 |
| N22S19 | 20 | rs3903703 | 12945963 | A | G | 0.32 | 0.33 | 0.046 | 2.31E-13 |
| N22S19 | 20 | rs4814173 | 12947532 | G | C | 0.32 | 0.33 | 0.046 | 2.31E-13 |
| N22S19 | 20 | rs3848744 | 12942649 | A | G | 0.32 | 0.33 | 0.046 | 2.84E-13 |
| N22S19 | 20 | rs3843765 | 12943737 | G | A | 0.32 | 0.33 | 0.046 | 3.52E-13 |
| N22S19 | 20 | rs3848745 | 12944067 | G | A | 0.32 | 0.33 | 0.046 | 3.52E-13 |
| N22S19 | 20 | rs6041735 | 12940649 | T | C | 0.32 | 0.33 | 0.046 | 3.97E-13 |
| N22S19 | 20 | rs4813102 | 12947883 | A | T | 0.37 | 0.30 | 0.045 | 9.50E-12 |
| N23S20 | 20 | rs1321940 | 12959885 | A | G | 0.37 | 0.32 | 0.048 | 4.62E-11 |
| N23S20 | 20 | rs364585 | 12962718 | A | G | 0.37 | 0.32 | 0.048 | 4.62E-11 |
| N23S20 | 20 | rs168622 | 12966089 | T | G | 0.37 | 0.32 | 0.048 | 4.62E-11 |
| N23S20 | 20 | rs680379 | 12969400 | A | G | 0.37 | 0.32 | 0.048 | 5.14E-11 |
| N23S20 | 20 | rs686548 | 12973521 | A | T | 0.37 | 0.32 | 0.048 | 5.14E-11 |
| N23S20 | 20 | rs438568 | 12958687 | A | G | 0.37 | 0.31 | 0.048 | 7.07E-11 |
| N23S20 | 20 | rs4814175 | 12959094 | A | T | 0.37 | 0.31 | 0.048 | 2.07E-10 |
| N23S20 | 20 | rs4814176 | 12959398 | T | C | 0.37 | 0.31 | 0.048 | 2.07E-10 |
| N23S20 | 20 | rs2327451 | 12953934 | C | A | 0.32 | 0.28 | 0.050 | 1.50E-08 |
| N23S20 | 20 | rs4508668 | 12955601 | T | C | 0.32 | 0.28 | 0.050 | 1.50E-08 |
| N23S20 | 20 | rs2327452 | 12952964 | A | C | 0.32 | 0.28 | 0.050 | 1.70E-08 |
| N23S20 | 20 | rs3903703 | 12945963 | A | G | 0.32 | 0.28 | 0.050 | 1.72E-08 |
| N23S20 | 20 | rs4814173 | 12947532 | G | C | 0.32 | 0.28 | 0.050 | 1.72E-08 |
| N23S20 | 20 | rs3848746 | 12950606 | A | G | 0.32 | 0.28 | 0.050 | 2.28E-08 |
| N23S20 | 20 | rs4813102 | 12947883 | A | T | 0.37 | 0.27 | 0.049 | 2.90E-08 |
| N24DS19 | 20 | rs1321940 | 12959885 | A | G | 0.37 | 0.44 | 0.047 | 3.14E-21 |
| N24DS19 | 20 | rs364585 | 12962718 | A | G | 0.37 | 0.44 | 0.047 | 3.14E-21 |
| N24DS19 | 20 | rs168622 | 12966089 | T | G | 0.37 | 0.44 | 0.047 | 3.14E-21 |
| N24DS19 | 20 | rs438568 | 12958687 | A | G | 0.36 | 0.44 | 0.047 | 4.18E-21 |
| N24DS19 | 20 | rs680379 | 12969400 | A | G | 0.37 | 0.43 | 0.046 | 1.13E-20 |
| N24DS19 | 20 | rs686548 | 12973521 | A | T | 0.37 | 0.43 | 0.046 | 1.13E-20 |
| N24DS19 | 20 | rs4814175 | 12959094 | A | T | 0.37 | 0.43 | 0.047 | 1.85E-20 |
| N24DS19 | 20 | rs4814176 | 12959398 | T | C | 0.37 | 0.43 | 0.047 | 1.85E-20 |
| N24DS19 | 20 | rs3848746 | 12950606 | A | G | 0.32 | 0.41 | 0.049 | 2.37E-17 |
| N24DS19 | 20 | rs2327451 | 12953934 | C | A | 0.32 | 0.40 | 0.049 | 1.11E-16 |
| N24DS19 | 20 | rs4508668 | 12955601 | T | C | 0.32 | 0.40 | 0.049 | 1.11E-16 |
| N24DS19 | 20 | rs3903703 | 12945963 | A | G | 0.32 | 0.40 | 0.049 | 1.29E-16 |
| N24DS19 | 20 | rs4814173 | 12947532 | G | C | 0.32 | 0.40 | 0.049 | 1.29E-16 |
| N24DS19 | 20 | rs2327452 | 12952964 | A | C | 0.32 | 0.40 | 0.049 | 1.40E-16 |
| N24DS19 | 20 | rs3843765 | 12943737 | G | A | 0.32 | 0.40 | 0.049 | 2.55E-16 |
| N24DS19 | 20 | rs3848745 | 12944067 | G | A | 0.32 | 0.40 | 0.049 | 2.55E-16 |
| N24DS19 | 20 | rs6041735 | 12940649 | T | C | 0.32 | 0.40 | 0.049 | 2.86E-16 |
| N24DS19 | 20 | rs3848744 | 12942649 | A | G | 0.32 | 0.40 | 0.049 | 3.21E-16 |
| N24DS19 | 20 | rs4813102 | 12947883 | A | T | 0.37 | 0.34 | 0.047 | 4.74E-13 |
| N24DS19 | 20 | rs6078854 | 12960153 | A | T | 0.40 | -0.30 | 0.046 | 4.65E-11 |
| N24DS19 | 20 | rs4544513 | 12954215 | T | C | 0.35 | -0.31 | 0.047 | 5.46E-11 |
| N24DS19 | 20 | rs6109637 | 12954804 | T | C | 0.35 | -0.31 | 0.047 | 5.46E-11 |
| N24DS19 | 20 | rs382003 | 12963171 | A | G | 0.30 | -0.29 | 0.049 | 1.59E-09 |
| N24DS19 | 20 | rs360539 | 12966440 | G | T | 0.30 | -0.29 | 0.048 | 2.57E-09 |
| N24DS19 | 20 | rs73079703 | 12941782 | T | C | 0.30 | -0.29 | 0.049 | 3.06E-09 |
| N24DS19 | 20 | rs8183164 | 12942600 | A | C | 0.30 | -0.29 | 0.049 | 3.06E-09 |
| N24DS19 | 20 | rs6131414 | 12945669 | A | G | 0.30 | -0.29 | 0.049 | 3.06E-09 |
| N24DS19 | 20 | rs7272107 | 12946328 | A | G | 0.30 | -0.29 | 0.049 | 3.06E-09 |
| N24DS19 | 20 | rs73079713 | 12947141 | T | C | 0.30 | -0.29 | 0.049 | 3.06E-09 |
| N24DS19 | 20 | rs6134734 | 12952640 | T | A | 0.30 | -0.29 | 0.049 | 4.20E-09 |
| N24DS19 | 20 | rs6109634 | 12953314 | G | A | 0.30 | -0.29 | 0.049 | 4.20E-09 |
| N24DS19 | 20 | rs3848748 | 12957587 | C | G | 0.29 | -0.29 | 0.049 | 4.56E-09 |
| N24DS19 | 20 | rs3848749 | 12962089 | C | T | 0.29 | -0.29 | 0.049 | 4.56E-09 |
| N24DS19 | 20 | rs6131417 | 12967751 | A | G | 0.29 | -0.28 | 0.049 | 7.26E-09 |
| N24DS19 | 20 | rs6134740 | 12968649 | C | G | 0.29 | -0.28 | 0.049 | 7.26E-09 |
| N24DS19 | 20 | rs6134741 | 12970842 | A | G | 0.29 | -0.28 | 0.049 | 7.26E-09 |
| N24DS19 | 20 | rs59131252 | 12975257 | A | C | 0.29 | -0.27 | 0.049 | 2.02E-08 |
| N24DS20 | 20 | rs680379 | 12969400 | A | G | 0.37 | 0.30 | 0.047 | 1.48E-10 |
| N24DS20 | 20 | rs686548 | 12973521 | A | T | 0.37 | 0.30 | 0.047 | 1.48E-10 |
| N24DS20 | 20 | rs1321940 | 12959885 | A | G | 0.37 | 0.30 | 0.047 | 1.79E-10 |
| N24DS20 | 20 | rs364585 | 12962718 | A | G | 0.37 | 0.30 | 0.047 | 1.79E-10 |
| N24DS20 | 20 | rs168622 | 12966089 | T | G | 0.37 | 0.30 | 0.047 | 1.79E-10 |
| N24DS20 | 20 | rs438568 | 12958687 | A | G | 0.37 | 0.30 | 0.047 | 2.62E-10 |
| N24DS20 | 20 | rs4814175 | 12959094 | A | T | 0.37 | 0.30 | 0.047 | 2.89E-10 |
| N24DS20 | 20 | rs4814176 | 12959398 | T | C | 0.37 | 0.30 | 0.047 | 2.89E-10 |
| N24DS20 | 20 | rs3848746 | 12950606 | A | G | 0.32 | 0.28 | 0.050 | 1.88E-08 |
| N24DS20 | 20 | rs2327451 | 12953934 | C | A | 0.32 | 0.28 | 0.050 | 2.34E-08 |
| N24DS20 | 20 | rs4508668 | 12955601 | T | C | 0.32 | 0.28 | 0.050 | 2.34E-08 |
| N24DS20 | 20 | rs2327452 | 12952964 | A | C | 0.32 | 0.27 | 0.050 | 2.81E-08 |
| N24DS20 | 20 | rs3903703 | 12945963 | A | G | 0.32 | 0.27 | 0.050 | 3.01E-08 |
| N24DS20 | 20 | rs4814173 | 12947532 | G | C | 0.32 | 0.27 | 0.050 | 3.01E-08 |
| N24S16 | 14 | rs7160525 | 64232220 | A | G | 0.14 | 0.36 | 0.058 | 5.67E-10 |
| N24S16 | 14 | rs17101394 | 64232386 | A | G | 0.14 | 0.36 | 0.058 | 5.67E-10 |
| N24S16 | 14 | rs8008068 | 64233717 | G | A | 0.14 | 0.36 | 0.058 | 5.67E-10 |
| N24S16 | 14 | rs8008070 | 64233720 | T | A | 0.14 | 0.36 | 0.058 | 5.67E-10 |
| N24S16 | 14 | rs8012828 | 64233980 | T | C | 0.14 | 0.36 | 0.058 | 5.67E-10 |
| N24S16 | 14 | rs34609767 | 64234034 | G | T | 0.14 | 0.36 | 0.058 | 5.67E-10 |
| N24S16 | 14 | rs4902243 | 64234243 | G | A | 0.14 | 0.36 | 0.058 | 5.67E-10 |
| N24S16 | 14 | **rs7157785** | 64235556 | T | G | 0.14 | 0.36 | 0.058 | 5.67E-10 |
| N24S16 | 14 | rs34817779 | 64236003 | T | C | 0.14 | 0.36 | 0.058 | 5.67E-10 |
| N24S16 | 14 | rs35372182 | 64236157 | G | A | 0.14 | 0.36 | 0.058 | 5.67E-10 |
| N24S16 | 14 | rs12897637 | 64239351 | C | T | 0.14 | 0.36 | 0.058 | 5.67E-10 |
| N24S16 | 14 | rs12878001 | 64239629 | G | T | 0.14 | 0.36 | 0.058 | 5.67E-10 |
| N24S16 | 20 | rs3848746 | 12950606 | A | G | 0.32 | 0.27 | 0.045 | 1.63E-09 |
| N24S16 | 20 | rs2327452 | 12952964 | A | C | 0.32 | 0.27 | 0.045 | 2.33E-09 |
| N24S16 | 20 | rs2327451 | 12953934 | C | A | 0.32 | 0.26 | 0.045 | 3.63E-09 |
| N24S16 | 20 | rs4508668 | 12955601 | T | C | 0.32 | 0.26 | 0.045 | 3.63E-09 |
| N24S16 | 20 | rs3903703 | 12945963 | A | G | 0.32 | 0.26 | 0.045 | 3.92E-09 |
| N24S16 | 20 | rs4814173 | 12947532 | G | C | 0.32 | 0.26 | 0.045 | 3.92E-09 |
| N24S16 | 20 | rs6041735 | 12940649 | T | C | 0.32 | 0.26 | 0.045 | 4.35E-09 |
| N24S16 | 20 | rs438568 | 12958687 | A | G | 0.36 | 0.25 | 0.043 | 5.41E-09 |
| N24S16 | 20 | rs680379 | 12969400 | A | G | 0.37 | 0.25 | 0.043 | 5.76E-09 |
| N24S16 | 20 | rs686548 | 12973521 | A | T | 0.37 | 0.25 | 0.043 | 5.76E-09 |
| N24S16 | 20 | rs3848744 | 12942649 | A | G | 0.32 | 0.26 | 0.045 | 5.83E-09 |
| N24S16 | 20 | rs1321940 | 12959885 | A | G | 0.37 | 0.25 | 0.043 | 8.37E-09 |
| N24S16 | 20 | rs364585 | 12962718 | A | G | 0.37 | 0.25 | 0.043 | 8.37E-09 |
| N24S16 | 20 | rs168622 | 12966089 | T | G | 0.37 | 0.25 | 0.043 | 8.37E-09 |
| N24S16 | 20 | rs3843765 | 12943737 | G | A | 0.32 | 0.26 | 0.045 | 8.98E-09 |
| N24S16 | 20 | rs3848745 | 12944067 | G | A | 0.32 | 0.26 | 0.045 | 8.98E-09 |
| N24S16 | 20 | rs4814175 | 12959094 | A | T | 0.37 | 0.24 | 0.043 | 2.58E-08 |
| N24S16 | 20 | rs4814176 | 12959398 | T | C | 0.37 | 0.24 | 0.043 | 2.58E-08 |
| N24S19 | 20 | rs438568 | 12958687 | A | G | 0.37 | 0.47 | 0.043 | 1.00E-27 |
| N24S19 | 20 | rs1321940 | 12959885 | A | G | 0.37 | 0.47 | 0.043 | 1.92E-27 |
| N24S19 | 20 | rs364585 | 12962718 | A | G | 0.37 | 0.47 | 0.043 | 1.92E-27 |
| N24S19 | 20 | rs168622 | 12966089 | T | G | 0.37 | 0.47 | 0.043 | 1.92E-27 |
| N24S19 | 20 | rs680379 | 12969400 | A | G | 0.37 | 0.46 | 0.043 | 4.82E-27 |
| N24S19 | 20 | rs686548 | 12973521 | A | T | 0.37 | 0.46 | 0.043 | 4.82E-27 |
| N24S19 | 20 | rs4814175 | 12959094 | A | T | 0.37 | 0.45 | 0.043 | 5.64E-26 |
| N24S19 | 20 | rs4814176 | 12959398 | T | C | 0.37 | 0.45 | 0.043 | 5.64E-26 |
| N24S19 | 20 | rs3848746 | 12950606 | A | G | 0.33 | 0.43 | 0.045 | 5.81E-22 |
| N24S19 | 20 | rs2327452 | 12952964 | A | C | 0.32 | 0.43 | 0.045 | 7.61E-22 |
| N24S19 | 20 | rs2327451 | 12953934 | C | A | 0.32 | 0.43 | 0.045 | 1.05E-21 |
| N24S19 | 20 | rs4508668 | 12955601 | T | C | 0.32 | 0.43 | 0.045 | 1.05E-21 |
| N24S19 | 20 | rs3903703 | 12945963 | A | G | 0.32 | 0.43 | 0.045 | 1.32E-21 |
| N24S19 | 20 | rs4814173 | 12947532 | G | C | 0.32 | 0.43 | 0.045 | 1.32E-21 |
| N24S19 | 20 | rs6041735 | 12940649 | T | C | 0.32 | 0.42 | 0.045 | 9.75E-21 |
| N24S19 | 20 | rs3848744 | 12942649 | A | G | 0.32 | 0.42 | 0.045 | 1.04E-20 |
| N24S19 | 20 | rs3843765 | 12943737 | G | A | 0.32 | 0.42 | 0.045 | 1.42E-20 |
| N24S19 | 20 | rs3848745 | 12944067 | G | A | 0.32 | 0.42 | 0.045 | 1.42E-20 |
| N24S19 | 20 | rs4813102 | 12947883 | A | T | 0.37 | 0.38 | 0.044 | 5.60E-18 |
| N24S19 | 20 | rs608994 | 12980885 | G | A | 0.31 | 0.30 | 0.044 | 2.92E-11 |
| N24S19 | 20 | rs6078854 | 12960153 | A | T | 0.40 | -0.28 | 0.042 | 4.29E-11 |
| N24S19 | 20 | rs3910136 | 12962261 | A | T | 0.34 | -0.26 | 0.044 | 4.61E-09 |
| N24S19 | 20 | rs6041755 | 12973617 | T | C | 0.33 | -0.25 | 0.044 | 1.07E-08 |
| N24S19 | 20 | rs4544513 | 12954215 | T | C | 0.35 | -0.25 | 0.043 | 1.61E-08 |
| N24S19 | 20 | rs6109637 | 12954804 | T | C | 0.35 | -0.25 | 0.043 | 1.61E-08 |
| N24S19 | 20 | rs3848754 | 12971345 | C | T | 0.28 | -0.26 | 0.047 | 3.51E-08 |
| N24S19 | 20 | rs3848755 | 12971437 | C | T | 0.28 | -0.26 | 0.047 | 3.51E-08 |
| N24S19 | 20 | rs13037956 | 12974302 | A | C | 0.28 | -0.26 | 0.047 | 3.51E-08 |
| N24S19 | 20 | rs6074538 | 12974493 | T | C | 0.28 | -0.26 | 0.047 | 3.51E-08 |
| N24S19 | 20 | rs6078866 | 12974567 | G | A | 0.28 | -0.26 | 0.047 | 3.51E-08 |
| N24S19 | 20 | rs6074539 | 12974665 | A | G | 0.28 | -0.26 | 0.047 | 3.51E-08 |
| N24S20 | 20 | rs680379 | 12969400 | A | G | 0.37 | 0.38 | 0.049 | 9.86E-15 |
| N24S20 | 20 | rs686548 | 12973521 | A | T | 0.37 | 0.38 | 0.049 | 9.86E-15 |
| N24S20 | 20 | rs1321940 | 12959885 | A | G | 0.37 | 0.38 | 0.049 | 1.20E-14 |
| N24S20 | 20 | rs364585 | 12962718 | A | G | 0.37 | 0.38 | 0.049 | 1.20E-14 |
| N24S20 | 20 | rs168622 | 12966089 | T | G | 0.37 | 0.38 | 0.049 | 1.20E-14 |
| N24S20 | 20 | rs438568 | 12958687 | A | G | 0.37 | 0.38 | 0.049 | 2.59E-14 |
| N24S20 | 20 | rs4814175 | 12959094 | A | T | 0.37 | 0.37 | 0.050 | 5.79E-14 |
| N24S20 | 20 | rs4814176 | 12959398 | T | C | 0.37 | 0.37 | 0.050 | 5.79E-14 |
| N24S20 | 20 | rs2327451 | 12953934 | C | A | 0.33 | 0.34 | 0.052 | 4.27E-11 |
| N24S20 | 20 | rs4508668 | 12955601 | T | C | 0.33 | 0.34 | 0.052 | 4.27E-11 |
| N24S20 | 20 | rs3903703 | 12945963 | A | G | 0.33 | 0.34 | 0.052 | 5.46E-11 |
| N24S20 | 20 | rs4814173 | 12947532 | G | C | 0.33 | 0.34 | 0.052 | 5.46E-11 |
| N24S20 | 20 | rs2327452 | 12952964 | A | C | 0.33 | 0.34 | 0.052 | 5.60E-11 |
| N24S20 | 20 | rs3848746 | 12950606 | A | G | 0.33 | 0.34 | 0.052 | 6.22E-11 |
| N24S20 | 20 | rs3843765 | 12943737 | G | A | 0.32 | 0.33 | 0.052 | 1.22E-10 |
| N24S20 | 20 | rs3848745 | 12944067 | G | A | 0.32 | 0.33 | 0.052 | 1.22E-10 |
| N24S20 | 20 | rs3848744 | 12942649 | A | G | 0.32 | 0.33 | 0.052 | 1.59E-10 |
| N24S20 | 20 | rs6041735 | 12940649 | T | C | 0.32 | 0.33 | 0.052 | 3.08E-10 |
| N24S20 | 20 | rs4813102 | 12947883 | A | T | 0.38 | 0.30 | 0.050 | 1.44E-09 |
| N25S20 | 20 | rs680379 | 12969400 | A | G | 0.37 | 0.29 | 0.048 | 9.18E-10 |
| N25S20 | 20 | rs686548 | 12973521 | A | T | 0.37 | 0.29 | 0.048 | 9.18E-10 |
| N25S20 | 20 | rs1321940 | 12959885 | A | G | 0.37 | 0.29 | 0.048 | 1.10E-09 |
| N25S20 | 20 | rs364585 | 12962718 | A | G | 0.37 | 0.29 | 0.048 | 1.10E-09 |
| N25S20 | 20 | rs168622 | 12966089 | T | G | 0.37 | 0.29 | 0.048 | 1.10E-09 |
| N25S20 | 20 | rs438568 | 12958687 | A | G | 0.36 | 0.29 | 0.048 | 1.19E-09 |
| N25S20 | 20 | rs4814175 | 12959094 | A | T | 0.37 | 0.29 | 0.048 | 2.08E-09 |
| N25S20 | 20 | rs4814176 | 12959398 | T | C | 0.37 | 0.29 | 0.048 | 2.08E-09 |
| N26S19 | 20 | rs438568 | 12958687 | A | G | 0.36 | 0.33 | 0.045 | 3.40E-13 |
| N26S19 | 20 | rs1321940 | 12959885 | A | G | 0.36 | 0.32 | 0.045 | 3.98E-13 |
| N26S19 | 20 | rs364585 | 12962718 | A | G | 0.36 | 0.32 | 0.045 | 3.98E-13 |
| N26S19 | 20 | rs168622 | 12966089 | T | G | 0.36 | 0.32 | 0.045 | 3.98E-13 |
| N26S19 | 20 | rs680379 | 12969400 | A | G | 0.37 | 0.32 | 0.045 | 7.18E-13 |
| N26S19 | 20 | rs686548 | 12973521 | A | T | 0.37 | 0.32 | 0.045 | 7.18E-13 |
| N26S19 | 20 | rs4814175 | 12959094 | A | T | 0.37 | 0.32 | 0.045 | 1.14E-12 |
| N26S19 | 20 | rs4814176 | 12959398 | T | C | 0.37 | 0.32 | 0.045 | 1.14E-12 |
| N26S19 | 6 | rs6940658 | 14238511 | C | G | 0.09 | 0.48 | 0.075 | 1.10E-10 |
| N26S19 | 6 | rs4333409 | 14240330 | A | C | 0.09 | 0.48 | 0.075 | 1.10E-10 |
| N26S19 | 6 | rs2039310 | 14250304 | A | G | 0.09 | 0.48 | 0.075 | 1.10E-10 |
| N26S19 | 6 | rs9382948 | 14251751 | A | G | 0.09 | 0.48 | 0.075 | 1.10E-10 |
| N26S19 | 6 | rs6910045 | 14255808 | G | A | 0.09 | 0.48 | 0.075 | 1.10E-10 |
| N26S19 | 6 | rs9367828 | 14236554 | G | A | 0.09 | 0.48 | 0.075 | 2.00E-10 |
| N26S19 | 6 | **rs9370735** | 14244517 | T | G | 0.09 | 0.48 | 0.075 | 2.00E-10 |
| N26S19 | 6 | rs6940973 | 14238655 | C | T | 0.09 | 0.47 | 0.076 | 8.61E-10 |
| N26S19 | 6 | rs1537152 | 14240435 | C | A | 0.09 | 0.47 | 0.076 | 8.61E-10 |
| N26S19 | 6 | **rs1537151** | 14240594 | C | A | 0.09 | 0.47 | 0.076 | 8.61E-10 |
| N26S19 | 6 | rs9396477 | 14234971 | C | G | 0.08 | 0.44 | 0.078 | 1.54E-08 |
| N26S19 | 20 | rs3848746 | 12950606 | A | G | 0.32 | 0.26 | 0.047 | 1.97E-08 |
| N26S19 | 20 | rs6041735 | 12940649 | T | C | 0.32 | 0.26 | 0.047 | 2.00E-08 |
| N26S19 | 6 | rs12208698 | 14237070 | G | T | 0.07 | 0.46 | 0.082 | 2.07E-08 |
| N26S19 | 6 | rs12190393 | 14239216 | A | G | 0.07 | 0.46 | 0.082 | 2.07E-08 |
| N26S19 | 6 | rs12212956 | 14240006 | G | A | 0.07 | 0.46 | 0.082 | 2.07E-08 |
| N26S19 | 6 | **rs12207359** | 14244274 | T | G | 0.07 | 0.46 | 0.082 | 2.07E-08 |
| N26S19 | 6 | rs75762794 | 14245458 | C | G | 0.07 | 0.46 | 0.082 | 2.07E-08 |
| N26S19 | 6 | rs2876349 | 14246531 | T | C | 0.07 | 0.46 | 0.082 | 2.07E-08 |
| N26S19 | 6 | rs12213267 | 14247608 | C | G | 0.07 | 0.46 | 0.082 | 2.07E-08 |
| N26S19 | 6 | rs115366574 | 14250710 | C | T | 0.07 | 0.46 | 0.082 | 2.07E-08 |
| N26S19 | 6 | rs79263173 | 14253258 | T | A | 0.07 | 0.46 | 0.082 | 2.07E-08 |
| N26S19 | 20 | rs2327452 | 12952964 | A | C | 0.32 | 0.26 | 0.047 | 2.29E-08 |
| N26S19 | 20 | rs3848744 | 12942649 | A | G | 0.32 | 0.26 | 0.047 | 2.55E-08 |
| N26S19 | 20 | rs2327451 | 12953934 | C | A | 0.32 | 0.26 | 0.047 | 2.92E-08 |
| N26S19 | 20 | rs4508668 | 12955601 | T | C | 0.32 | 0.26 | 0.047 | 2.92E-08 |
| N26S19 | 20 | rs3843765 | 12943737 | G | A | 0.32 | 0.26 | 0.047 | 3.25E-08 |
| N26S19 | 20 | rs3848745 | 12944067 | G | A | 0.32 | 0.26 | 0.047 | 3.25E-08 |
| N26S19 | 20 | rs3903703 | 12945963 | A | G | 0.32 | 0.26 | 0.047 | 3.31E-08 |
| N26S19 | 20 | rs4814173 | 12947532 | G | C | 0.32 | 0.26 | 0.047 | 3.31E-08 |
| N24S19ratio | 1 | rs4653568 | 224396329 | A | G | 0.35 | 0.29 | 0.046 | 2.36E-10 |
| N24S19ratio | 1 | rs4654000 | 224396487 | C | T | 0.35 | 0.29 | 0.046 | 2.36E-10 |
| N24S19ratio | 1 | rs9793489 | 224397459 | T | A | 0.35 | 0.29 | 0.046 | 2.36E-10 |
| N24S19ratio | 1 | rs2011117 | 224398639 | C | T | 0.35 | 0.29 | 0.046 | 2.36E-10 |
| N24S19ratio | 1 | rs908801 | 224398817 | T | C | 0.35 | 0.29 | 0.046 | 2.36E-10 |
| N24S19ratio | 1 | rs6681673 | 224398910 | T | C | 0.35 | 0.29 | 0.046 | 2.36E-10 |
| N24S19ratio | 1 | rs4654003 | 224399383 | T | C | 0.35 | 0.29 | 0.046 | 2.36E-10 |
| N24S19ratio | 1 | rs6682292 | 224400175 | A | G | 0.35 | 0.29 | 0.046 | 2.36E-10 |
| N24S19ratio | 1 | rs12038372 | 224363881 | T | C | 0.41 | 0.26 | 0.045 | 1.30E-08 |
| N24S19ratio | 1 | rs6426143 | 224318875 | G | A | 0.40 | 0.25 | 0.045 | 2.15E-08 |
| N24S19ratio | 1 | rs6682551 | 224352258 | C | T | 0.39 | 0.25 | 0.045 | 2.42E-08 |
| N24S19ratio | 1 | **rs10799505** | 224351023 | T | C | 0.40 | 0.25 | 0.045 | 3.24E-08 |
| N24S19ratio | 1 | rs10916403 | 224352551 | C | A | 0.40 | 0.25 | 0.045 | 3.24E-08 |
| N24S19ratio | 1 | rs55664906 | 224355682 | A | C | 0.40 | 0.25 | 0.045 | 3.24E-08 |
| N24S19ratio | 1 | rs12076788 | 224355701 | C | A | 0.40 | 0.25 | 0.045 | 3.24E-08 |
| N24S19ratio | 1 | rs4653563 | 224314776 | A | C | 0.40 | 0.25 | 0.045 | 3.68E-08 |
| N24S19ratio | 1 | rs4653564 | 224314814 | C | T | 0.40 | 0.25 | 0.045 | 3.68E-08 |
| N24S19ratio | 1 | rs6426139 | 224315128 | A | G | 0.40 | 0.25 | 0.045 | 3.68E-08 |
| N24S19ratio | 1 | rs10753454 | 224316813 | C | T | 0.40 | 0.25 | 0.045 | 3.68E-08 |
| N24S19ratio | 1 | rs7542293 | 224316881 | A | G | 0.40 | 0.25 | 0.045 | 3.68E-08 |
| N24S19ratio | 1 | rs7542474 | 224317043 | A | C | 0.40 | 0.25 | 0.045 | 3.68E-08 |
| N24S19ratio | 1 | rs6698041 | 224317551 | G | A | 0.40 | 0.25 | 0.045 | 3.68E-08 |
| N24S19ratio | 1 | rs7519433 | 224319003 | T | C | 0.40 | 0.25 | 0.045 | 3.68E-08 |
| N24S19ratio | 1 | rs12730611 | 224319825 | G | C | 0.40 | 0.25 | 0.045 | 3.68E-08 |
| N24S19ratio | 1 | rs2014782 | 224321492 | T | C | 0.40 | 0.25 | 0.045 | 3.68E-08 |
| N24S19ratio | 1 | rs869945 | 224321637 | G | A | 0.40 | 0.25 | 0.045 | 3.68E-08 |
| N24S19ratio | 1 | rs6691405 | 224322879 | A | G | 0.40 | 0.25 | 0.045 | 3.68E-08 |
| N24S19ratio | 1 | rs6685783 | 224322942 | T | C | 0.40 | 0.25 | 0.045 | 3.68E-08 |
| N24S19ratio | 1 | rs10916355 | 224323548 | T | C | 0.40 | 0.25 | 0.045 | 3.68E-08 |
| N24S19ratio | 1 | rs66506223 | 224325436 | C | G | 0.40 | 0.25 | 0.045 | 3.68E-08 |
| N24S19ratio | 1 | rs61827699 | 224325460 | G | A | 0.40 | 0.25 | 0.045 | 3.68E-08 |
| N24S19ratio | 1 | rs10916367 | 224329969 | G | A | 0.40 | 0.25 | 0.045 | 4.33E-08 |
| N24S19ratio | 1 | rs4653986 | 224330818 | A | G | 0.40 | 0.25 | 0.045 | 4.33E-08 |
| N24S19ratio | 1 | rs7518839 | 224331779 | G | A | 0.40 | 0.25 | 0.045 | 4.33E-08 |
| N24S19ratio | 1 | rs13374070 | 224332078 | A | G | 0.40 | 0.25 | 0.045 | 4.33E-08 |
| N24S19ratio | 1 | rs55736782 | 224334072 | G | A | 0.40 | 0.25 | 0.045 | 4.33E-08 |
| N24S19ratio | 1 | rs997297 | 224335492 | A | T | 0.40 | 0.25 | 0.045 | 4.33E-08 |
| N24S19ratio | 1 | rs997296 | 224335497 | C | G | 0.40 | 0.25 | 0.045 | 4.33E-08 |
| N24S19ratio | 1 | rs7531891 | 224335956 | A | G | 0.40 | 0.25 | 0.045 | 4.33E-08 |
| N24S19ratio | 1 | rs1492694 | 224337153 | G | A | 0.40 | 0.25 | 0.045 | 4.33E-08 |
| N24S19ratio | 1 | rs4653991 | 224337208 | G | A | 0.40 | 0.25 | 0.045 | 4.33E-08 |
| N24S19ratio | 1 | rs7546235 | 224338017 | T | C | 0.40 | 0.25 | 0.045 | 4.33E-08 |
| N24S19ratio | 1 | rs10916371 | 224338162 | A | G | 0.40 | 0.25 | 0.045 | 4.33E-08 |
| N24S19ratio | 1 | rs7524705 | 224339000 | A | G | 0.40 | 0.25 | 0.045 | 4.33E-08 |
| N24S19ratio | 1 | **rs1826421** | 224339447 | A | G | 0.40 | 0.25 | 0.045 | 4.33E-08 |
| N24S19ratio | 1 | rs12563153 | 224339647 | A | G | 0.40 | 0.25 | 0.045 | 4.33E-08 |
| N24S19ratio | 1 | rs8328 | 224346959 | G | A | 0.40 | 0.25 | 0.045 | 4.33E-08 |
| N24S19ratio | 1 | rs7526252 | 224348818 | T | C | 0.40 | 0.25 | 0.045 | 4.33E-08 |
| S19Sum | 20 | rs438568 | 12958687 | A | G | 0.37 | 0.48 | 0.043 | 4.46E-29 |
| S19Sum | 20 | rs1321940 | 12959885 | A | G | 0.37 | 0.48 | 0.043 | 6.33E-29 |
| S19Sum | 20 | rs364585 | 12962718 | A | G | 0.37 | 0.48 | 0.043 | 6.33E-29 |
| S19Sum | 20 | rs168622 | 12966089 | T | G | 0.37 | 0.48 | 0.043 | 6.33E-29 |
| S19Sum | 20 | rs680379 | 12969400 | A | G | 0.37 | 0.48 | 0.043 | 2.19E-28 |
| S19Sum | 20 | rs686548 | 12973521 | A | T | 0.37 | 0.48 | 0.043 | 2.19E-28 |
| S19Sum | 20 | rs4814175 | 12959094 | A | T | 0.37 | 0.47 | 0.043 | 2.37E-27 |
| S19Sum | 20 | rs4814176 | 12959398 | T | C | 0.37 | 0.47 | 0.043 | 2.37E-27 |
| S19Sum | 20 | rs3848746 | 12950606 | A | G | 0.33 | 0.44 | 0.045 | 1.11E-22 |
| S19Sum | 20 | rs2327452 | 12952964 | A | C | 0.32 | 0.44 | 0.045 | 1.56E-22 |
| S19Sum | 20 | rs2327451 | 12953934 | C | A | 0.32 | 0.44 | 0.045 | 1.95E-22 |
| S19Sum | 20 | rs4508668 | 12955601 | T | C | 0.32 | 0.44 | 0.045 | 1.95E-22 |
| S19Sum | 20 | rs3903703 | 12945963 | A | G | 0.32 | 0.44 | 0.045 | 2.68E-22 |
| S19Sum | 20 | rs4814173 | 12947532 | G | C | 0.32 | 0.44 | 0.045 | 2.68E-22 |
| S19Sum | 20 | rs3848744 | 12942649 | A | G | 0.32 | 0.43 | 0.045 | 2.12E-21 |
| S19Sum | 20 | rs3843765 | 12943737 | G | A | 0.32 | 0.43 | 0.045 | 2.64E-21 |
| S19Sum | 20 | rs3848745 | 12944067 | G | A | 0.32 | 0.43 | 0.045 | 2.64E-21 |
| S19Sum | 20 | rs6041735 | 12940649 | T | C | 0.32 | 0.43 | 0.045 | 3.28E-21 |
| S19Sum | 20 | rs4813102 | 12947883 | A | T | 0.37 | 0.39 | 0.044 | 7.47E-19 |
| S19Sum | 20 | rs6078854 | 12960153 | A | T | 0.40 | -0.30 | 0.042 | 3.38E-12 |
| S19Sum | 20 | rs608994 | 12980885 | G | A | 0.31 | 0.30 | 0.045 | 1.70E-11 |
| S19Sum | 20 | rs4544513 | 12954215 | T | C | 0.35 | -0.26 | 0.044 | 1.79E-09 |
| S19Sum | 20 | rs6109637 | 12954804 | T | C | 0.35 | -0.26 | 0.044 | 1.79E-09 |
| S19Sum | 20 | rs382003 | 12963171 | A | G | 0.30 | -0.26 | 0.045 | 6.06E-09 |
| S19Sum | 20 | rs360539 | 12966440 | G | T | 0.30 | -0.26 | 0.045 | 7.23E-09 |
| S19Sum | 20 | rs3910136 | 12962261 | A | T | 0.34 | -0.25 | 0.044 | 8.71E-09 |
| S19Sum | 20 | rs73079703 | 12941782 | T | C | 0.30 | -0.25 | 0.045 | 2.40E-08 |
| S19Sum | 20 | rs8183164 | 12942600 | A | C | 0.30 | -0.25 | 0.045 | 2.40E-08 |
| S19Sum | 20 | rs6131414 | 12945669 | A | G | 0.30 | -0.25 | 0.045 | 2.40E-08 |
| S19Sum | 20 | rs7272107 | 12946328 | A | G | 0.30 | -0.25 | 0.045 | 2.40E-08 |
| S19Sum | 20 | rs73079713 | 12947141 | T | C | 0.30 | -0.25 | 0.045 | 2.40E-08 |
| S19Sum | 20 | rs6041755 | 12973617 | T | C | 0.33 | -0.25 | 0.044 | 2.42E-08 |
| S19Sum | 20 | rs6134734 | 12952640 | T | A | 0.30 | -0.25 | 0.045 | 3.94E-08 |
| S19Sum | 20 | rs6109634 | 12953314 | G | A | 0.30 | -0.25 | 0.045 | 3.94E-08 |
| S19Sum | 20 | rs3848748 | 12957587 | C | G | 0.29 | -0.25 | 0.045 | 4.26E-08 |
| S19Sum | 20 | rs3848749 | 12962089 | C | T | 0.29 | -0.25 | 0.045 | 4.26E-08 |
| S19Sum | 20 | rs3848754 | 12971345 | C | T | 0.28 | -0.26 | 0.047 | 4.90E-08 |
| S19Sum | 20 | rs3848755 | 12971437 | C | T | 0.28 | -0.26 | 0.047 | 4.90E-08 |
| S19Sum | 20 | rs13037956 | 12974302 | A | C | 0.28 | -0.26 | 0.047 | 4.90E-08 |
| S19Sum | 20 | rs6074538 | 12974493 | T | C | 0.28 | -0.26 | 0.047 | 4.90E-08 |
| S19Sum | 20 | rs6078866 | 12974567 | G | A | 0.28 | -0.26 | 0.047 | 4.90E-08 |
| S19Sum | 20 | rs6074539 | 12974665 | A | G | 0.28 | -0.26 | 0.047 | 4.90E-08 |
| S20Sum | 20 | rs680379 | 12969400 | A | G | 0.37 | 0.40 | 0.050 | 5.17E-16 |
| S20Sum | 20 | rs686548 | 12973521 | A | T | 0.37 | 0.40 | 0.050 | 5.17E-16 |
| S20Sum | 20 | rs1321940 | 12959885 | A | G | 0.37 | 0.40 | 0.050 | 5.94E-16 |
| S20Sum | 20 | rs364585 | 12962718 | A | G | 0.37 | 0.40 | 0.050 | 5.94E-16 |
| S20Sum | 20 | rs168622 | 12966089 | T | G | 0.37 | 0.40 | 0.050 | 5.94E-16 |
| S20Sum | 20 | rs438568 | 12958687 | A | G | 0.37 | 0.40 | 0.050 | 1.12E-15 |
| S20Sum | 20 | rs4814175 | 12959094 | A | T | 0.37 | 0.39 | 0.050 | 3.56E-15 |
| S20Sum | 20 | rs4814176 | 12959398 | T | C | 0.37 | 0.39 | 0.050 | 3.56E-15 |
| S20Sum | 20 | rs2327451 | 12953934 | C | A | 0.33 | 0.35 | 0.052 | 1.31E-11 |
| S20Sum | 20 | rs4508668 | 12955601 | T | C | 0.33 | 0.35 | 0.052 | 1.31E-11 |
| S20Sum | 20 | rs2327452 | 12952964 | A | C | 0.32 | 0.35 | 0.052 | 1.47E-11 |
| S20Sum | 20 | rs3903703 | 12945963 | A | G | 0.32 | 0.35 | 0.052 | 1.74E-11 |
| S20Sum | 20 | rs4814173 | 12947532 | G | C | 0.32 | 0.35 | 0.052 | 1.74E-11 |
| S20Sum | 20 | rs3848746 | 12950606 | A | G | 0.33 | 0.35 | 0.052 | 1.77E-11 |
| S20Sum | 20 | rs3843765 | 12943737 | G | A | 0.32 | 0.34 | 0.052 | 4.04E-11 |
| S20Sum | 20 | rs3848745 | 12944067 | G | A | 0.32 | 0.34 | 0.052 | 4.04E-11 |
| S20Sum | 20 | rs3848744 | 12942649 | A | G | 0.32 | 0.34 | 0.052 | 4.52E-11 |
| S20Sum | 20 | rs6041735 | 12940649 | T | C | 0.32 | 0.34 | 0.052 | 8.35E-11 |
| S20Sum | 20 | rs4813102 | 12947883 | A | T | 0.38 | 0.32 | 0.051 | 3.91E-10 |

Table 11A: Ensembl and GTEx summary of the significant SNPs identified by GWAS association for ceramides and related sphingolipid species.
Description of the SNPs using the Ensembl API Client and summary of the information on eQTL status as identified using the GTEX browser.

| SNP ID | Gene Name | Gene Type | Variant Allele | Consequence Terms | GTEx | Liver or whole blood? | Other |
| --- | --- | --- | --- | --- | --- | --- | --- |
| rs438568 | LINC01723 | lncRNA | G | intron | SPTLC3 | Liver | X |
| rs1321940 | LINC01723 | lncRNA | G | intron | SPTLC3 | Liver | ISM1 (pancreas) |
| rs364585 | LINC01723 | lncRNA | G | downstream | SPTLC3 | Liver | X |
| rs168622 | LINC01723 | lncRNA | G | intron | SPTLC3 | Liver | X |
| rs680379 | LINC01723 | lncRNA | G | intron | SPTLC3 | Liver | X |
| rs686548 | LINC01723 | lncRNA | T | intron | SPTLC3 | Liver | ISM1 (pancreas) |
| rs4814175 | LINC01723 | lncRNA | T | intron | SPTLC3 | Liver | X |
| rs4814176 | LINC01723 | lncRNA | C | intron | SPTLC3 | Liver | X |
| rs2327452 | LINC01723 | lncRNA | C | intron | SPTLC3 | Liver | X |
| rs3848746 | LINC01723 | lncRNA | G | intron | SPTLC3 | Liver | X |
| rs2327451 | LINC01723 | lncRNA | A | intron | SPTLC3 | Liver | X |
| rs4508668 | LINC01723 | lncRNA | C | intron | SPTLC3 | Liver | X |
| rs3903703 | LINC01723 | lncRNA | G | intron | SPTLC3 | Liver | X |
| rs4814173 | LINC01723 | lncRNA | A | intron | SPTLC3 | Liver | X |
| rs3848744 | LINC01723 | lncRNA | G | intron | SPTLC3 | Liver | X |
| rs3843765 | LINC01723 | lncRNA | A | intron | SPTLC3 | Liver | X |
| rs3848745 | LINC01723 | lncRNA | A | intron | SPTLC3 | Liver | X |
| rs6041735 | LINC01723 | lncRNA | C | intron | SPTLC3 | Liver | X |
| rs4813102 | LINC01723 | lncRNA | T | intron | SPTLC3 | Liver | X |
| rs6078854 | LINC01723 | lncRNA | A | downstream | SPTLC3 | X | X |
| rs4544513 | LINC01723 | lncRNA | T | intron | SPTLC3 | Liver | X |
| rs6109637 | LINC01723 | lncRNA | T | intron | SPTLC3 | Liver | X |
| rs382003 | LINC01723 | lncRNA | A | downstream | SPTLC3 | Liver | X |
| rs360539 | LINC01723 | lncRNA | C | intron | SPTLC3 | Liver | X |
| rs73079703 | LINC01723 | lncRNA | T | intron | X | X | X |
| rs8183164 | LINC01723 | lncRNA | A | intron | X | X | X |
| rs6131414 | LINC01723 | lncRNA | A | intron | X | X | X |
| rs7272107 | LINC01723 | lncRNA | A | intron | X | X | X |
| rs73079713 | LINC01723 | lncRNA | T | intron | X | X | X |
| rs6134734 | LINC01723 | lncRNA | T | intron | X | X | X |
| rs6109634 | LINC01723 | lncRNA | G | intron | X | X | X |
| rs3848748 | LINC01723 | lncRNA | C | intron | X | X | X |
| rs3848749 | LINC01723 | lncRNA | C | downstream | X | X | X |
| rs6131417 | LINC01723 | lncRNA | A | intron | X | X | X |
| rs6134740 | LINC01723 | lncRNA | C | intron | X | X | X |
| rs6134741 | LINC01723 | lncRNA | A | intron | X | X | X |
| rs59131252 | LINC01723 | lncRNA | A | intron | X | X | X |
| rs7160525 | AL161670.1 | pseudogene | A | downstream | X | X | X |
| rs17101394 | AL161670.1 | pseudogene | A | downstream | X | X | X |
| rs8008068 | AL161670.1 | pseudogene | G | downstream | X | X | X |
| rs8008070 | AL161670.1 | pseudogene | T | downstream | X | X | X |
| rs8012828 | Intergenic | Intergenic | T | intergenic | X | X | X |
| rs34609767 | Intergenic | Intergenic | G | intergenic | X | X | X |
| rs4902243 | Intergenic | Intergenic | G | intergenic | X | X | X |
| rs7157785 | Intergenic | Intergenic | T | intergenic | X | X | X |
| rs34817779 | Intergenic | Intergenic | T | intergenic | X | X | X |
| rs35372182 | Intergenic | Intergenic | G | intergenic | X | X | X |
| rs12897637 | Intergenic | Intergenic | C | intergenic | X | X | X |
| rs12878001 | Intergenic | Intergenic | G | intergenic | X | X | X |
| rs608994 | LINC01723 | lncRNA | A | intron | SPTLC3 | Liver | X |
| rs3910136 | LINC01723 | lncRNA | A | downstream | NA | NA | NA |
| rs6041755 | LINC01723 | lncRNA | T | intron | SPTLC3 | X | X |
| rs3848754 | LINC01723 | lncRNA | C | intron | SPTLC3 | X | X |
| rs3848755 | LINC01723 | lncRNA | C | intron | X | X | X |
| rs13037956 | LINC01723 | lncRNA | A | intron | SPTLC3 | X | X |
| rs6074538 | LINC01723 | lncRNA | T | intron | SPTLC3 | X | X |
| rs6078866 | LINC01723 | lncRNA | G | intron | SPTLC3 | X | X |
| rs6074539 | LINC01723 | lncRNA | A | intron | X | X | X |
| rs6940658 | Intergenic | Intergenic | G | intergenic | X | X | X |
| rs4333409 | Intergenic | Intergenic | C | intergenic | X | X | X |
| rs2039310 | Intergenic | Intergenic | C | intergenic | X | X | X |
| rs9382948 | Intergenic | Intergenic | G | intergenic | X | X | X |
| rs6910045 | Intergenic | Intergenic | A | intergenic | RP3-500L14.2 | X | X |
| rs9367828 | AL353152.1 | lncRNA | A | upstream | X | X | X |
| rs9370735 | Intergenic | Intergenic | G | intergenic | X | X | X |
| rs6940973 | Intergenic | Intergenic | T | intergenic | X | X | X |
| rs1537152 | Intergenic | Intergenic | A | intergenic | X | X | X |
| rs1537151 | Intergenic | Intergenic | A | intergenic | X | X | X |
| rs9396477 | AL353152.1 | lncRNA | G | upstream | X | X | X |
| rs12208698 | Intergenic | Intergenic | A | intergenic | X | X | X |
| rs12190393 | Intergenic | Intergenic | A | intergenic | X | X | X |
| rs12212956 | Intergenic | Intergenic | G | intergenic | X | X | X |
| rs12207359 | Intergenic | Intergenic | T | intergenic | X | X | X |
| rs75762794 | Intergenic | Intergenic | C | intergenic | X | X | X |
| rs2876349 | Intergenic | Intergenic | A | intergenic | X | X | X |
| rs12213267 | Intergenic | Intergenic | C | intergenic | X | X | X |
| rs115366574 | Intergenic | Intergenic | C | intergenic | X | X | X |
| rs79263173 | Intergenic | Intergenic | T | intergenic | X | X | X |
| rs4653568 | AC092809.2 | lncRNA | G | downstream | FBXO28 | blood | DEGS1, RP11-365O16.3, CAPN8 |
| rs4654000 | AC092809.2 | lncRNA | G | exon | FBXO28 | blood | DEGS1, RP11-365O16.3, CAPN8, GTP2IP20 |
| rs9793489 | AC092809.2 | lncRNA | A | intron | FBXO28 | blood | DEGS1, RP11-365O16.3, CAPN8, GTP2IP20 |
| rs2011117 | AC092809.2 | lncRNA | T | intron | FBXO28 | blood | DEGS1, RP11-365O16.3, CAPN8, GTP2IP20 |
| rs908801 | AC092809.2 | lncRNA | A | intron | FBXO28 | blood | DEGS1, RP11-365O16.3, CAPN8, GTP2IP20 |
| rs6681673 | AC092809.2 | lncRNA | C | intron | FBXO28 | blood | DEGS1, RP11-365O16.3, CAPN8, GTP2IP20 |
| rs4654003 | AC092809.2 | lncRNA | C | intron | FBXO28 | blood | DEGS1, RP11-365O16.3, CAPN8, GTP2IP20 |
| rs6682292 | AC092809.2 | lncRNA | G | intron | FBXO28 | blood | DEGS1, RP11-365O16.3, CAPN8, GTP2IP20 |
| rs12038372 | DEGS1 | protein_coding | G | intron | FBXO28 | blood | DEGS1, RP11-365O16.3, CAPN8 |
| rs6426143 | FBXO28 | protein_coding | A | intron | FBXO28 | blood | DEGS1, RP11-365O16.3, CAPN8 |
| rs6682551 | FBXO28 | protein_coding | C | downstream | FBXO28 | blood | RP11-365O16.3 |
| rs10799505 | FBXO28 | protein_coding | T | downstream | FBXO28 | blood | DEGS1, RP11-365O16.3, CAPN8, CNIH3 |
| rs10916403 | FBXO28 | protein_coding | C | downstream | FBXO28 | blood | DEGS1, RP11-365O16.3, CAPN8, CNIH3 |
| rs55664906 | Intergenic | Intergenic | A | intergenic | FBXO28 | blood | DEGS1, RP11-365O16.3, CAPN8 |
| rs12076788 | Intergenic | Intergenic | A | intergenic | FBXO28 | blood | DEGS1, RP11-365O16.3, CAPN8 |
| rs4653563 | FBXO28 | protein_coding | C | intron | FBXO28 | blood | DEGS1, RP11-365O16.3, CAPN8 |
| rs4653564 | FBXO28 | protein_coding | T | intron | FBXO28 | blood | DEGS1, RP11-365O16.3, CAPN8 |
| rs6426139 | FBXO28 | protein_coding | G | intron | FBXO28 | blood | DEGS1, RP11-365O16.3, CAPN8 |
| rs10753454 | FBXO28 | protein_coding | G | intron | FBXO28 | blood | DEGS1, RP11-365O16.3, CAPN8 |
| rs7542293 | FBXO28 | protein_coding | G | intron | FBXO28 | blood | DEGS1, RP11-365O16.3, CAPN8 |
| rs7542474 | FBXO28 | protein_coding | C | intron | FBXO28 | blood | DEGS1, RP11-365O16.3, CAPN8 |
| rs6698041 | FBXO28 | protein_coding | A | intron | FBXO28 | blood | DEGS1, RP11-365O16.3, CAPN8 |
| rs7519433 | FBXO28 | protein_coding | T | intron | FBXO28 | blood | RP11-365O16.3 |
| rs12730611 | FBXO28 | protein_coding | C | intron | FBXO28 | blood | DEGS1, RP11-365O16.3, CAPN8 |
| rs2014782 | FBXO28 | protein_coding | T | intron | FBXO28 | blood | DEGS1, RP11-365O16.3, CAPN8 |
| rs869945 | FBXO28 | protein_coding | G | intron | FBXO28 | blood | DEGS1, RP11-365O16.3, CAPN8 |
| rs6691405 | FBXO28 | protein_coding | G | intron | FBXO28 | blood | DEGS1, RP11-365O16.3, CAPN8 |
| rs6685783 | FBXO28 | protein_coding | C | intron | FBXO28 | blood | DEGS1, RP11-365O16.3, CAPN8 |
| rs10916355 | FBXO28 | protein_coding | T | intron | FBXO28 | blood | DEGS1, RP11-365O16.3, CAPN8 |
| rs66506223 | FBXO28 | protein_coding | G | intron | FBXO28 | blood | DEGS1, RP11-365O16.3, CAPN8 |
| rs61827699 | FBXO28 | protein_coding | A | intron | FBXO28 | blood | DEGS1, RP11-365O16.3, CAPN8 |
| rs10916367 | FBXO28 | protein_coding | A | intron | FBXO28 | blood | DEGS1, RP11-365O16.3, CAPN8 |
| rs4653986 | FBXO28 | protein_coding | G | intron | FBXO28 | blood | DEGS1, RP11-365O16.3, CAPN8 |
| rs7518839 | FBXO28 | protein_coding | A | intron | FBXO28 | blood | DEGS1, RP11-365O16.3, CAPN8 |
| rs13374070 | FBXO28 | protein_coding | A | intron | FBXO28 | blood | DEGS1, RP11-365O16.3, CAPN8 |
| rs55736782 | FBXO28 | protein_coding | G | intron | FBXO28 | blood | DEGS1, RP11-365O16.3, CAPN8 |
| rs997297 | FBXO28 | protein_coding | T | intron | FBXO28 | blood | DEGS1, RP11-365O16.3, CAPN8 |
| rs997296 | FBXO28 | protein_coding | A | intron | FBXO28 | blood | DEGS1, RP11-365O16.3, CAPN8 |
| rs7531891 | FBXO28 | protein_coding | A | intron | FBXO28 | blood | DEGS1, RP11-365O16.3, CAPN8 |
| rs1492694 | FBXO28 | protein_coding | G | intron | FBXO28 | blood | DEGS1, RP11-365O16.3, CAPN8 |
| rs4653991 | FBXO28 | protein_coding | A | intron | FBXO28 | blood | DEGS1, RP11-365O16.3, CAPN8 |
| rs7546235 | FBXO28 | protein_coding | T | intron | FBXO28 | blood | DEGS1, RP11-365O16.3, CAPN8 |
| rs10916371 | FBXO28 | protein_coding | G | intron | FBXO28 | blood | DEGS1, RP11-365O16.3, CAPN8 |
| rs7524705 | FBXO28 | protein_coding | A | intron | FBXO28 | blood | DEGS1, RP11-365O16.3, CAPN8 |
| rs1826421 | FBXO28 | protein_coding | G | intron | FBXO28 | blood | DEGS1, RP11-365O16.3, CAPN8 |
| rs12563153 | FBXO28 | protein_coding | A | intron | FBXO28 | blood | DEGS1, RP11-365O16.3, CAPN8 |
| rs8328 | FBXO28 | protein_coding | A | 3_prime_UTR | FBXO28 | blood | DEGS1, RP11-365O16.3, CAPN8 |
| rs7526252 | FBXO28 | protein_coding | C | 3_prime_UTR | FBXO28 | blood | DEGS1, RP11-365O16.3, CAPN8 |

Table S11B: GWAS Catalog searches, Gene Atlas PheWAS, and UCSC Genome Browser summaries of the significant SNPs identified by GWAS association for ceramides and related sphingolipid species
The table depicts the results of GWAS Catalog searches, Gene Atlas PheWAS, and known CER metabolic enzymes genes identified at that locus using UCSC Genome Browser.

| **SNP ID** | **GWAS Catalog** | **Gene Atlas** | **UCSC** |
| --- | --- | --- | --- |
| rs438568 | X | X | NA |
| rs1321940 | X | X | NA |
| rs364585 | LDL cholesterol | X | NA |
| rs168622 | X | X | NA |
| rs680379 | SL levels, FA levels, Glycerophospholipid levels | X | NA |
| rs686548 | Serum metabolite ratios in chronic kidney disease | X | NA |
| rs4814175 | X | X | NA |
| rs4814176 | SL levels, blood metabolites | X | NA |
| rs2327452 | X | X | NA |
| rs3848746 | X | X | NA |
| rs2327451 | X | X | NA |
| rs4508668 | X | X | NA |
| rs3903703 | FA levels | X | NA |
| rs4814173 | X | X | NA |
| rs3848744 | X | X | NA |
| rs3843765 | X | X | NA |
| rs3848745 | X | X | NA |
| rs6041735 | X | X | NA |
| rs4813102 | X | X | NA |
| rs6078854 | X | X | NA |
| rs4544513 | X | X | NA |
| rs6109637 | X | X | NA |
| rs382003 | X | X | NA |
| rs360539 | X | X | NA |
| rs73079703 | X | X | NA |
| rs8183164 | X | X | NA |
| rs6131414 | X | X | NA |
| rs7272107 | X | X | NA |
| rs73079713 | X | X | NA |
| rs6134734 | X | X | NA |
| rs6109634 | X | X | NA |
| rs3848748 | X | X | NA |
| rs3848749 | X | X | NA |
| rs6131417 | X | X | NA |
| rs6134740 | X | X | NA |
| rs6134741 | X | X | NA |
| rs59131252 | X | X | NA |
| rs7160525 | Serum metabolite concentrations in chronic kidney disease | Mean platelet (thrombocyte) volume (P=3.2825e-29); Red blood cell (erythrocyte) distribution width (P=5.963e-14); Platelet count (P=5.5601e-13); High light scatter reticulocyte percentage (P=1.2661e-12); High light scatter reticulocyte count (P=6.6865e-11); Immature reticulocyte fraction (P=1.0614e-10); Reticulocyte percentage (P=1.9119e-08) | SGPP1 |
| rs17101394 | SL levels | Mean platelet (thrombocyte) volume (P=5.1305e-29); Red blood cell (erythrocyte) distribution width (P=4.9311e-14); Platelet count (P=6.397e-13); High light scatter reticulocyte percentage (P=3.2699e-12); High light scatter reticulocyte count (P=1.1051e-10); Immature reticulocyte fraction (P=2.0931e-10); Reticulocyte percentage (P=1.8191e-08) | SGPP1 |
| rs8008068 | Red cell distribution width | Mean platelet (thrombocyte) volume (P=4.7089e-29); Red blood cell (erythrocyte) distribution width (P=4.8152e-14); Platelet count (P=6.2582e-13); High light scatter reticulocyte percentage (P=1.866e-12); Immature reticulocyte fraction (P=5.3687e-11); High light scatter reticulocyte count (P=6.6258e-11); Reticulocyte percentage (P=2.0003e-08) | SGPP1 |
| rs8008070 | Serum metabolite ratios in chronic kidney disease | Mean platelet (thrombocyte) volume (P=2.8823e-29); Red blood cell (erythrocyte) distribution width (P=4.8987e-14); Platelet count (P=5.7716e-13); High light scatter reticulocyte percentage (P=1.7608e-12); Immature reticulocyte fraction (P=4.8545e-11); High light scatter reticulocyte count (P=6.1781e-11); Reticulocyte percentage (P=1.9361e-08) | SGPP1 |
| rs8012828 | X | Mean platelet (thrombocyte) volume (P=4.6195e-29); Red blood cell (erythrocyte) distribution width (P=5.5573e-14); Platelet count (P=7.3757e-13); High light scatter reticulocyte percentage (P=3.4938e-12); High light scatter reticulocyte count (P=1.2452e-10); Immature reticulocyte fraction (P=2.1356e-10); Reticulocyte percentage (P=1.6812e-08) | SGPP1 |
| rs34609767 | X | Mean platelet (thrombocyte) volume (P=3.1253e-29); Red blood cell (erythrocyte) distribution width (P=4.292e-14); Platelet count (P=5.6134e-13); High light scatter reticulocyte percentage (P=1.75e-12); High light scatter reticulocyte count (P=6.6063e-11); Immature reticulocyte fraction (P=6.8857e-11); Reticulocyte percentage (P=1.6786e-08) | SGPP1 |
| rs4902243 | Blood metabolite levels | Mean platelet (thrombocyte) volume (P=4.2606e-29); Red blood cell (erythrocyte) distribution width (P=2.0997e-14); Platelet count (P=6.7994e-13); High light scatter reticulocyte percentage (P=2.5783e-12); High light scatter reticulocyte count (P=9.5096e-11); Immature reticulocyte fraction (P=1.0739e-10); Reticulocyte percentage (P=2.0268e-08) | SGPP1 |
| rs7157785 | SL levels, blood metabolites, glycerophospholipids, total cholesterol [Hicks, shin, draisma, other] | Mean platelet (thrombocyte) volume (P=1.2992e-28); Red blood cell (erythrocyte) distribution width (P=3.5732e-13); Platelet count (P=9.5427e-13); High light scatter reticulocyte percentage (P=1.0509e-12); Immature reticulocyte fraction (P=5.289e-11); High light scatter reticulocyte count (P=5.6669e-11); Reticulocyte percentage (P=1.6633e-08) | SGPP1 |
| rs34817779 | X | Mean platelet (thrombocyte) volume (P=3.6761e-29); Red blood cell (erythrocyte) distribution width (P=3.6016e-14); Platelet count (P=4.2642e-13); High light scatter reticulocyte percentage (P=1.5459e-12); High light scatter reticulocyte count (P=6.0295e-11); Immature reticulocyte fraction (P=8.1684e-11); Reticulocyte percentage (P=1.5888e-08) | SGPP1 |
| rs35372182 | X | Mean platelet (thrombocyte) volume (P=4.1731e-29); Red blood cell (erythrocyte) distribution width (P=4.1885e-14); Platelet count (P=4.8832e-13); High light scatter reticulocyte percentage (P=2.0206e-12); High light scatter reticulocyte count (P=7.8142e-11); Immature reticulocyte fraction (P=8.981e-11); Reticulocyte percentage (P=1.7224e-08) | SGPP1 |
| rs12897637 | red blood cell distribution width | Mean platelet (thrombocyte) volume (P=9.1577e-29); Red blood cell (erythrocyte) distribution width (P=1.2123e-14); Platelet count (P=3.9706e-13); High light scatter reticulocyte percentage (P=1.2183e-12); High light scatter reticulocyte count (P=4.5639e-11); Immature reticulocyte fraction (P=9.9928e-11); Reticulocyte percentage (P=1.2307e-08) | SGPP1 |
| rs12878001 | X | Mean platelet (thrombocyte) volume (P=4.1485e-29); Red blood cell (erythrocyte) distribution width (P=1.9038e-14); Platelet count (P=2.7341e-13); High light scatter reticulocyte percentage (P=7.549e-13); High light scatter reticulocyte count (P=2.948e-11); Immature reticulocyte fraction (P=5.8129e-11); Reticulocyte percentage (P=1.1537e-08) | SGPP1 |
| rs608994 | X | X | NA |
| rs3910136 | X | X | NA |
| rs6041755 | X | X | NA |
| rs3848754 | X | X | NA |
| rs3848755 | X | X | NA |
| rs13037956 | X | X | NA |
| rs6074538 | X | X | NA |
| rs6078866 | X | X | NA |
| rs6074539 | X | X | NA |
| rs6940658 | X | X | upstream to CD83 |
| rs4333409 | X | X | upstream to CD83 |
| rs2039310 | X | X | upstream to CD83 |
| rs9382948 | X | X | upstream to CD83 |
| rs6910045 | X | X | upstream to CD83 |
| rs9367828 | X | X | upstream to CD83 |
| rs9370735 | X | X | upstream to CD83 |
| rs6940973 | X | X | upstream to CD83 |
| rs1537152 | X | X | upstream to CD83 |
| rs1537151 | X | X | upstream to CD83 |
| rs9396477 | X | X | upstream to CD83 |
| rs12208698 | X | X | upstream to CD83 |
| rs12190393 | X | X | upstream to CD83 |
| rs12212956 | X | X | upstream to CD83 |
| rs12207359 | X | X | upstream to CD83 |
| rs75762794 | X | X | upstream to CD83 |
| rs2876349 | X | X | upstream to CD83 |
| rs12213267 | X | X | upstream to CD83 |
| rs115366574 | X | X | upstream to CD83 |
| rs79263173 | X | X | upstream to CD83 |
| rs4653568 | X | Mean platelet (thrombocyte) volume (P=4.7652e-12) | DEGS1 |
| rs4654000 | X | Mean platelet (thrombocyte) volume (P=4.8955e-12) | DEGS1 |
| rs9793489 | X | Mean platelet (thrombocyte) volume (P=6.0912e-12) | DEGS1 |
| rs2011117 | X | Mean platelet (thrombocyte) volume (P=6.3718e-12) | DEGS1 |
| rs908801 | X | Mean platelet (thrombocyte) volume (P=6.3548e-12) | DEGS1 |
| rs6681673 | X | Mean platelet (thrombocyte) volume (P=7.1624e-12) | DEGS1 |
| rs4654003 | X | Mean platelet (thrombocyte) volume (P=6.6526e-12) | DEGS1 |
| rs6682292 | X | Mean platelet (thrombocyte) volume (P=7.1866e-12) | DEGS1 |
| rs12038372 | X | Mean platelet (thrombocyte) volume (P=3.3948e-18); Neutrophil count (P=1.1357e-09); White blood cell (leukocyte) count (P=3.1852e-09)). | DEGS1 |
| rs6426143 | X | Mean platelet (thrombocyte) volume (P=1.9455e-16); White blood cell (leukocyte) count (P=4.4764e-10); Neutrophil count (P=1.7302e-09); Red blood cell (erythrocyte) count (P=2.3883e-09)). | DEGS1 |
| rs6682551 | X | Mean platelet (thrombocyte) volume (P=1.409e-16); White blood cell (leukocyte) count (P=2.7024e-10); Neutrophil count (P=4.8023e-09); Red blood cell (erythrocyte) count (P=1.2395e-09)). | DEGS1 |
| rs10799505 | X | Mean platelet (thrombocyte) volume (P=1.513e-16); White blood cell (leukocyte) count (P=4.7039e-10); Red blood cell (erythrocyte) count (P=5.487e-10); Neutrophil count (P=6.0446e-10)). | DEGS1 |
| rs10916403 | X | Mean platelet (thrombocyte) volume (P=1.6163e-16); White blood cell (leukocyte) count (P=4.6079e-10); Red blood cell (erythrocyte) count (P=5.9676e-10); Neutrophil count (P=6.0795e-10); Mean sphered cell volume (P=9.3161e-08). | DEGS1 |
| rs55664906 | X | Mean platelet (thrombocyte) volume (P=5.3355e-16); Red blood cell (erythrocyte) count (P=5.5676e-10); White blood cell (leukocyte) count (P=1.6824e-09); Neutrophil count (P=2.3745e-09); Mean sphered cell volume (P=6.4397e-08). | DEGS1 |
| rs12076788 | X | Mean platelet (thrombocyte) volume (P=2.1908e-16); White blood cell (leukocyte) count (P=7.174e-10); Red blood cell (erythrocyte) count (P=1.3034e-09); Neutrophil count (P=1.6361e-09). | DEGS1 |
| rs4653563 | X | Mean platelet (thrombocyte) volume (P=9.2972e-17); White blood cell (leukocyte) count (P=4.6256e-10); Neutrophil count (P=1.5481e-09); Red blood cell (erythrocyte) count (P=2.0336e-09). | DEGS1 |
| rs4653564 | X | Mean platelet (thrombocyte) volume (P=8.4965e-17); White blood cell (leukocyte) count (P=4.9323e-10); Neutrophil count (P=1.5762e-09); Red blood cell (erythrocyte) count (P=1.8868e-09). | DEGS1 |
| rs6426139 | X | Mean platelet (thrombocyte) volume (P=8.978e-17); White blood cell (leukocyte) count (P=4.5862e-10); Neutrophil count (P=1.5587e-09); Red blood cell (erythrocyte) count (P=2.0271e-09). | DEGS1 |
| rs10753454 | X | Mean platelet (thrombocyte) volume (P=1.0592e-16); White blood cell (leukocyte) count (P=4.7645e-10); Neutrophil count (P=1.5892e-09); Red blood cell (erythrocyte) count (P=1.8429e-09). | DEGS1 |
| rs7542293 | X | Mean platelet (thrombocyte) volume (P=1.0092e-16); White blood cell (leukocyte) count (P=4.5228e-10); Neutrophil count (P=1.5183e-09); Red blood cell (erythrocyte) count (P=1.8065e-09). | DEGS1 |
| rs7542474 | X | Mean platelet (thrombocyte) volume (P=9.8213e-17); White blood cell (leukocyte) count (P=4.5324e-10); Neutrophil count (P=1.5489e-09); Red blood cell (erythrocyte) count (P=1.8863e-09). | DEGS1 |
| rs6698041 | X | Mean platelet (thrombocyte) volume (P=1.0606e-16); White blood cell (leukocyte) count (P=4.7114e-10); Neutrophil count (P=1.5776e-09); Red blood cell (erythrocyte) count (P=1.8254e-09). | DEGS1 |
| rs7519433 | X | Mean platelet (thrombocyte) volume (P=9.8438e-17); White blood cell (leukocyte) count (P=3.8904e-10); Neutrophil count (P=1.2884e-09); Red blood cell (erythrocyte) count (P=2.125e-09). | DEGS1 |
| rs12730611 | X | Mean platelet (thrombocyte) volume (P=1.0549e-16); White blood cell (leukocyte) count (P=4.5558e-10); Neutrophil count (P=1.5474e-09); Red blood cell (erythrocyte) count (P=1.7702e-09). | DEGS1 |
| rs2014782 | X | Mean platelet (thrombocyte) volume (P=1.037e-16); White blood cell (leukocyte) count (P=3.6376e-10); Neutrophil count (P=7.8676e-10); Red blood cell (erythrocyte) count (P=1.4462e-09). | DEGS1 |
| rs869945 | X | Mean platelet (thrombocyte) volume (P=1.0322e-16); White blood cell (leukocyte) count (P=3.5835e-10); Neutrophil count (P=7.7447e-10); Red blood cell (erythrocyte) count (P=1.4451e-09). | DEGS1 |
| rs6691405 | X | Mean platelet (thrombocyte) volume (P=8.4246e-17); White blood cell (leukocyte) count (P=4.2562e-10); Neutrophil count (P=1.469e-09); Red blood cell (erythrocyte) count (P=2.0154e-09). | DEGS1 |
| rs6685783 | X | Mean platelet (thrombocyte) volume (P=9.2798e-17); White blood cell (leukocyte) count (P=4.0593e-10); Neutrophil count (P=1.4225e-09); Red blood cell (erythrocyte) count (P=2.0518e-09). | DEGS1 |
| rs10916355 | X | Mean platelet (thrombocyte) volume (P=8.8465e-17); White blood cell (leukocyte) count (P=3.6897e-10); Neutrophil count (P=7.8979e-10); Red blood cell (erythrocyte) count (P=1.3939e-09). | DEGS1 |
| rs66506223 | X | Mean platelet (thrombocyte) volume (P=8.4007e-17); White blood cell (leukocyte) count (P=4.1628e-10); Neutrophil count (P=1.4926e-09); Red blood cell (erythrocyte) count (P=2.1213e-09). | DEGS1 |
| rs61827699 | X | Mean platelet (thrombocyte) volume (P=8.882e-17); White blood cell (leukocyte) count (P=3.9042e-10); Neutrophil count (P=1.3987e-09); Red blood cell (erythrocyte) count (P=2.1437e-09). | DEGS1 |
| rs10916367 | X | Mean platelet (thrombocyte) volume (P=5.0835e-17); White blood cell (leukocyte) count (P=3.555e-10); Neutrophil count (P=1.21e-09); Red blood cell (erythrocyte) count (P=1.8198e-09). | DEGS1 |
| rs4653986 | X | Mean platelet (thrombocyte) volume (P=4.6912e-17); White blood cell (leukocyte) count (P=3.513e-10); Neutrophil count (P=1.1833e-09); Red blood cell (erythrocyte) count (P=1.8124e-09). | DEGS1 |
| rs7518839 | X | Mean platelet (thrombocyte) volume (P=4.9221e-17); White blood cell (leukocyte) count (P=3.4999e-10); Neutrophil count (P=1.1542e-09); Red blood cell (erythrocyte) count (P=1.8229e-09). | DEGS1 |
| rs13374070 | X | Mean platelet (thrombocyte) volume (P=4.8718e-17); White blood cell (leukocyte) count (P=2.794e-10); Neutrophil count (P=6.354e-10); Red blood cell (erythrocyte) count (P=1.0861e-09). | DEGS1 |
| rs55736782 | X | Mean platelet (thrombocyte) volume (P=4.8498e-17); White blood cell (leukocyte) count (P=2.7262e-10); Neutrophil count (P=6.1833e-10); Red blood cell (erythrocyte) count (P=1.0713e-09). | DEGS1 |
| rs997297 | X | Mean platelet (thrombocyte) volume (P=3.7928e-17); White blood cell (leukocyte) count (P=3.6642e-10); Neutrophil count (P=1.3338e-09); Red blood cell (erythrocyte) count (P=1.8344e-09). | DEGS1 |
| rs997296 | X | Mean platelet (thrombocyte) volume (P=3.1302e-17); White blood cell (leukocyte) count (P=3.2187e-10); Neutrophil count (P=1.1007e-09); Red blood cell (erythrocyte) count (P=1.5943e-09). | DEGS1 |
| rs7531891 | X | Mean platelet (thrombocyte) volume (P=6.0753e-17); White blood cell (leukocyte) count (P=2.5459e-10); Neutrophil count (P=5.6997e-10); Red blood cell (erythrocyte) count (P=9.9016e-10). | DEGS1 |
| rs1492694 | X | Mean platelet (thrombocyte) volume (P=5.3205e-17); White blood cell (leukocyte) count (P=2.5298e-10); Neutrophil count (P=5.7417e-10); Red blood cell (erythrocyte) count (P=1.0494e-09). | DEGS1 |
| rs4653991 | X | Mean platelet (thrombocyte) volume (P=5.0426e-17); White blood cell (leukocyte) count (P=3.204e-10); Neutrophil count (P=1.0394e-09); Red blood cell (erythrocyte) count (P=1.7487e-09). | DEGS1 |
| rs7546235 | X | Mean platelet (thrombocyte) volume (P=5.2812e-17); White blood cell (leukocyte) count (P=2.4989e-10); Neutrophil count (P=5.7025e-10); Red blood cell (erythrocyte) count (P=1.0533e-09). | DEGS1 |
| rs10916371 | X | Mean platelet (thrombocyte) volume (P=4.6957e-17); White blood cell (leukocyte) count (P=3.03e-10); Neutrophil count (P=9.7973e-10); Red blood cell (erythrocyte) count (P=1.7318e-09). | DEGS1 |
| rs7524705 | X | Mean platelet (thrombocyte) volume (P=4.7556e-17); White blood cell (leukocyte) count (P=2.4762e-10); Neutrophil count (P=5.7017e-10); Red blood cell (erythrocyte) count (P=1.0728e-09). | DEGS1 |
| rs1826421 | X | Mean platelet (thrombocyte) volume (P=4.5592e-17); White blood cell (leukocyte) count (P=3.3766e-10); Neutrophil count (P=1.1157e-09); Red blood cell (erythrocyte) count (P=1.733e-09). | DEGS1 |
| rs12563153 | X | Mean platelet (thrombocyte) volume (P=4.8579e-17); White blood cell (leukocyte) count (P=2.6386e-10); Neutrophil count (P=6.0969e-10); Red blood cell (erythrocyte) count (P=1.0513e-09). | DEGS1 |
| rs8328 | X | Mean platelet (thrombocyte) volume (P=5.1273e-17); White blood cell (leukocyte) count (P=3.3169e-10); Neutrophil count (P=1.1078e-09); Red blood cell (erythrocyte) count (P=1.7271e-09). | DEGS1 |
| rs7526252 | X | Mean platelet (thrombocyte) volume (P=4.3542e-17); White blood cell (leukocyte) count (P=3.5044e-10); Neutrophil count (P=1.2329e-09); Red blood cell (erythrocyte) count (P=1.7174e-09). | DEGS1 |

**Table S12. Significant results from the two-sample Mendelian randomisation analysis.**

The table depicts the significant association of CER[N(24)S(16)] after adjustment for 71 tests as exposures to multiple blood cell count outcomes via 2SMR analysis. More information on the study (via id) and outcome can be found in Appendix Table 0.14. The beta and standard error (SE) values describe the relationship between the ceramide traits and blood cell phenotypes.

| **ID** | **Outcome** | **Exposure** | **beta** | **SE** | **Padj** |
| --- | --- | --- | --- | --- | --- |
| 1247 | Mean platelet volume | CER[N(24)S(16)] | -0.08 | 0.014 | 8.13E-07 |
| 1251 | Platelet count | CER[N(24)S(16)] | 0.06 | 0.014 | 3.35E-03 |
| 1259 | High light scatter reticulocyte count | CER[N(24)S(16)] | -0.06 | 0.014 | 8.68E-04 |
| 1260 | High light scatter reticulocyte percentage of red cells | CER[N(24)S(16)] | -0.07 | 0.014 | 7.22E-05 |
| 1267 | Reticulocyte fraction of red cells | CER[N(24)S(16)] | -0.06 | 0.014 | 4.40E-04 |
| 1269 | Red cell distribution width | CER[N(24)S(16)] | 0.08 | 0.013 | 1.62E-08 |
| 1270 | Reticulocyte count | CER[N(24)S(16)] | -0.05 | 0.014 | 1.80E-02 |
| 1275 | Lymphocyte counts | CER[N(24)S(16)] | 0.05 | 0.014 | 2.54E-02 |
| 1276 | Immature fraction of reticulocytes | CER[N(24)S(16)] | -0.05 | 0.013 | 6.94E-03 |

**Supplemental Figures**

**A**

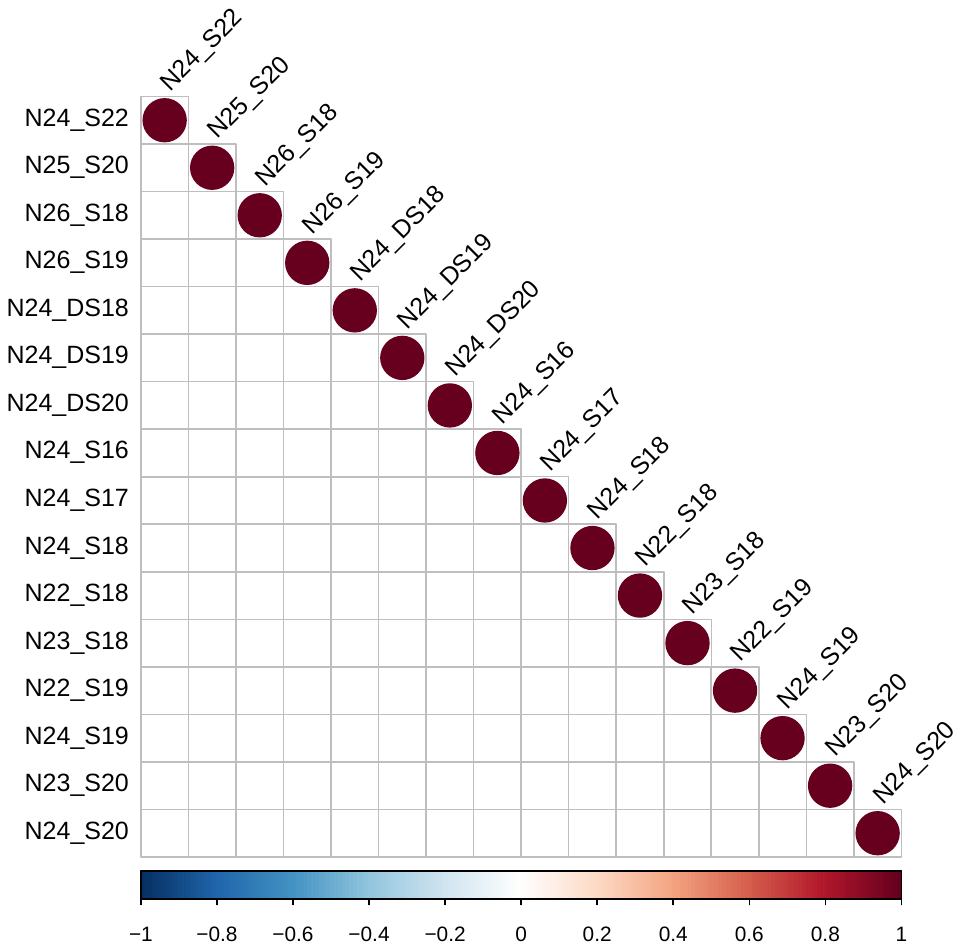

**B**

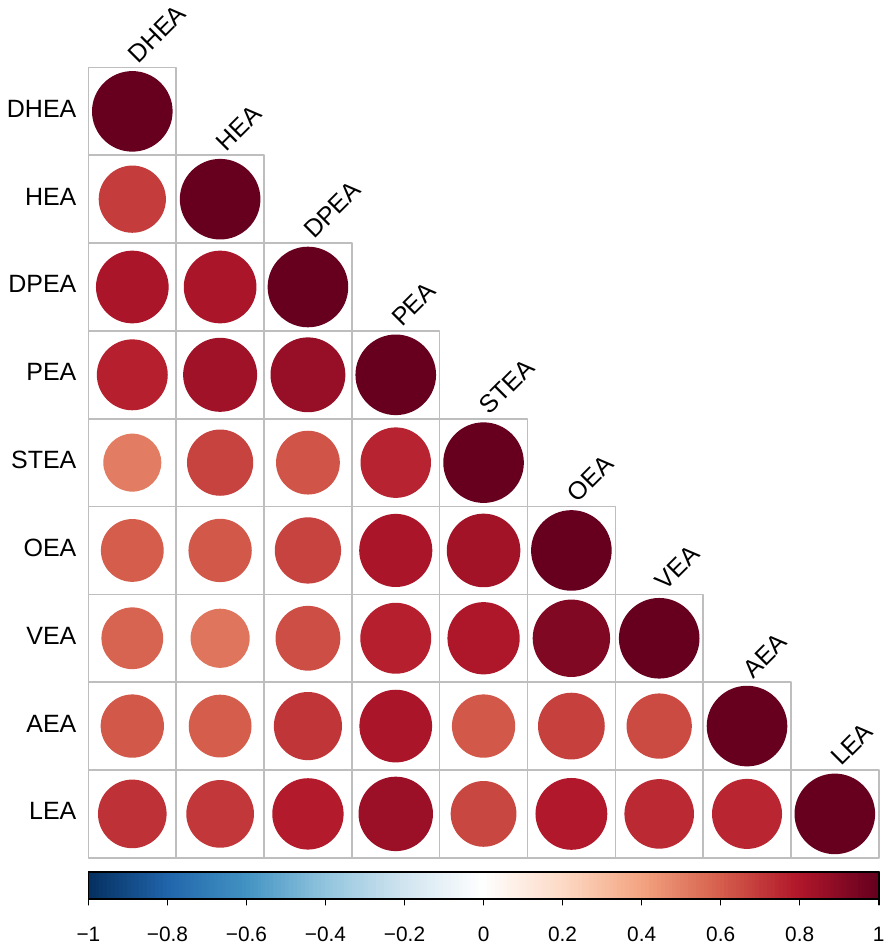

**Figure S1.** Intra-correlation analysis of ceramides, sphingoid bases, and related sphingolipids, and *N*-acyl ethanolamines.

The figure depicts the assessment of the relatedness within the two classes of lipids (A: CER, B: NAE) for the 999 plasma samples studied. The tool rquery.cormat was used in R, which takes into account strength of relationship (correlation coefficient; depicted as a scale of colours) and P-value (size of circle produced). The correlation was completed on the covariate-adjusted, standardised residuals used for genetic analyses.

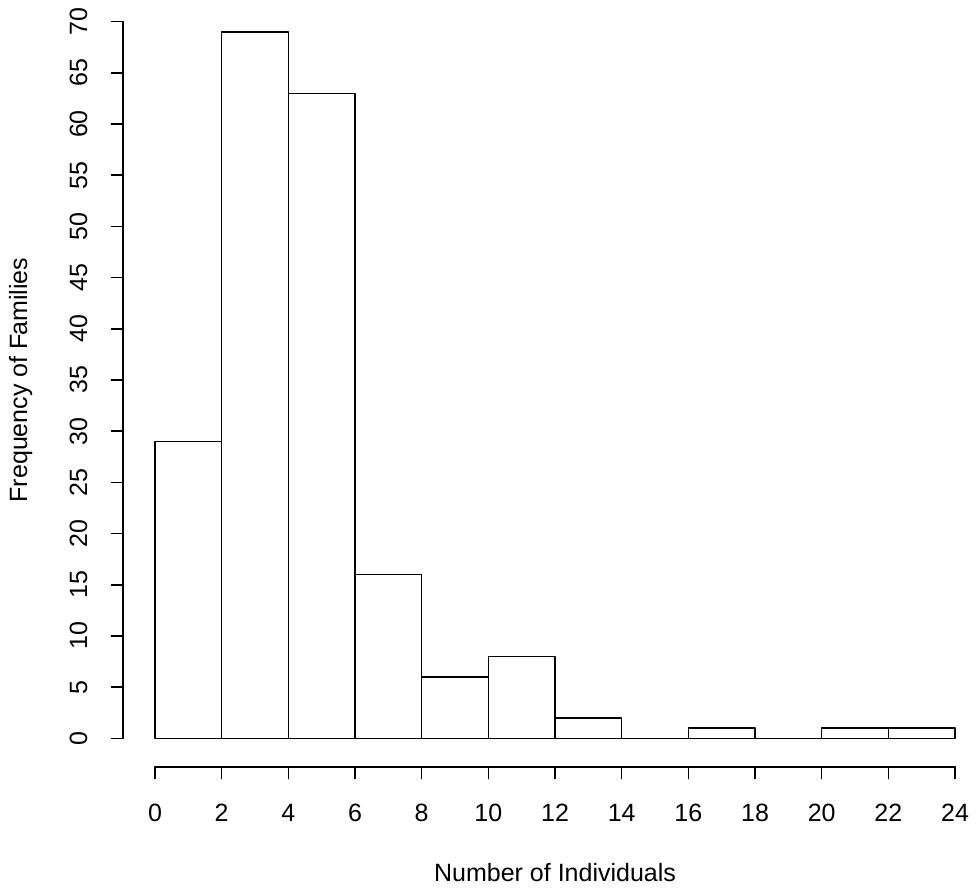

**Figure S2.** The frequency of individuals in each collected family that were analysed in this lipidomics study.

The graph depicts the range of individuals (1-24) with plasma available for lipidomics analyses. 999 individuals were assessed from 196 families. The mean number of individuals from each family assessed for mediator lipidomics was 5.

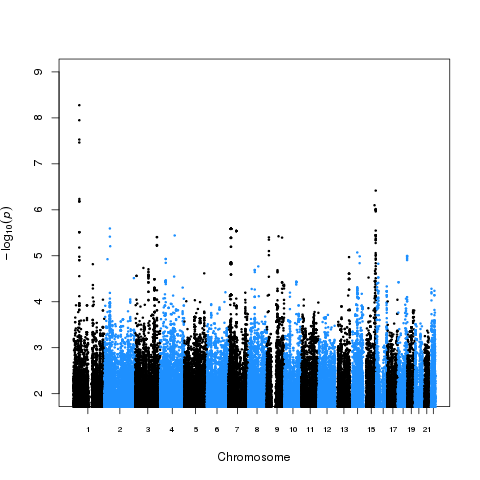

**Figure S3.** Manhattan plot of an example NAE species (PEA) associated at GWAS with SNPs in *FAAH* on chromosome 1.

The figure shows a Manhattan plot of GWAS results for PEA lipid species highlighting significant SNPs at chromosome 1 at the fatty acid amide hydrolase gene (*FAAH*) in 993 plasma samples.

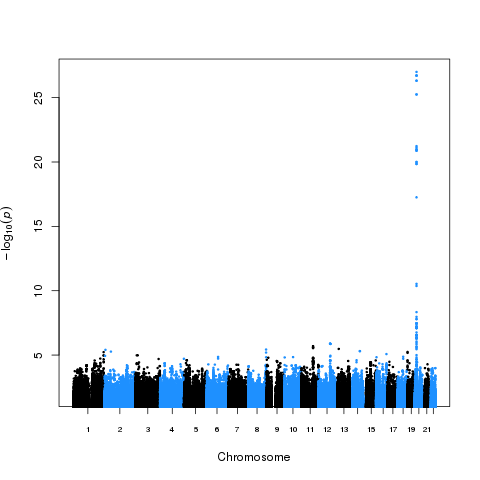

**Figure S4.** Manhattan plot of an example ceramide species N(24)S(19) associated at GWAS with SNPs in *SPTLC3* on chromosome 20.

The figure shows a Manhattan plot of GWAS results for CER[N(24)S(19)], highlighting significant SNPs at chromosome 20 at the serine palmitoyltransferase gene (*SPTLC3*) in 991 plasma samples.
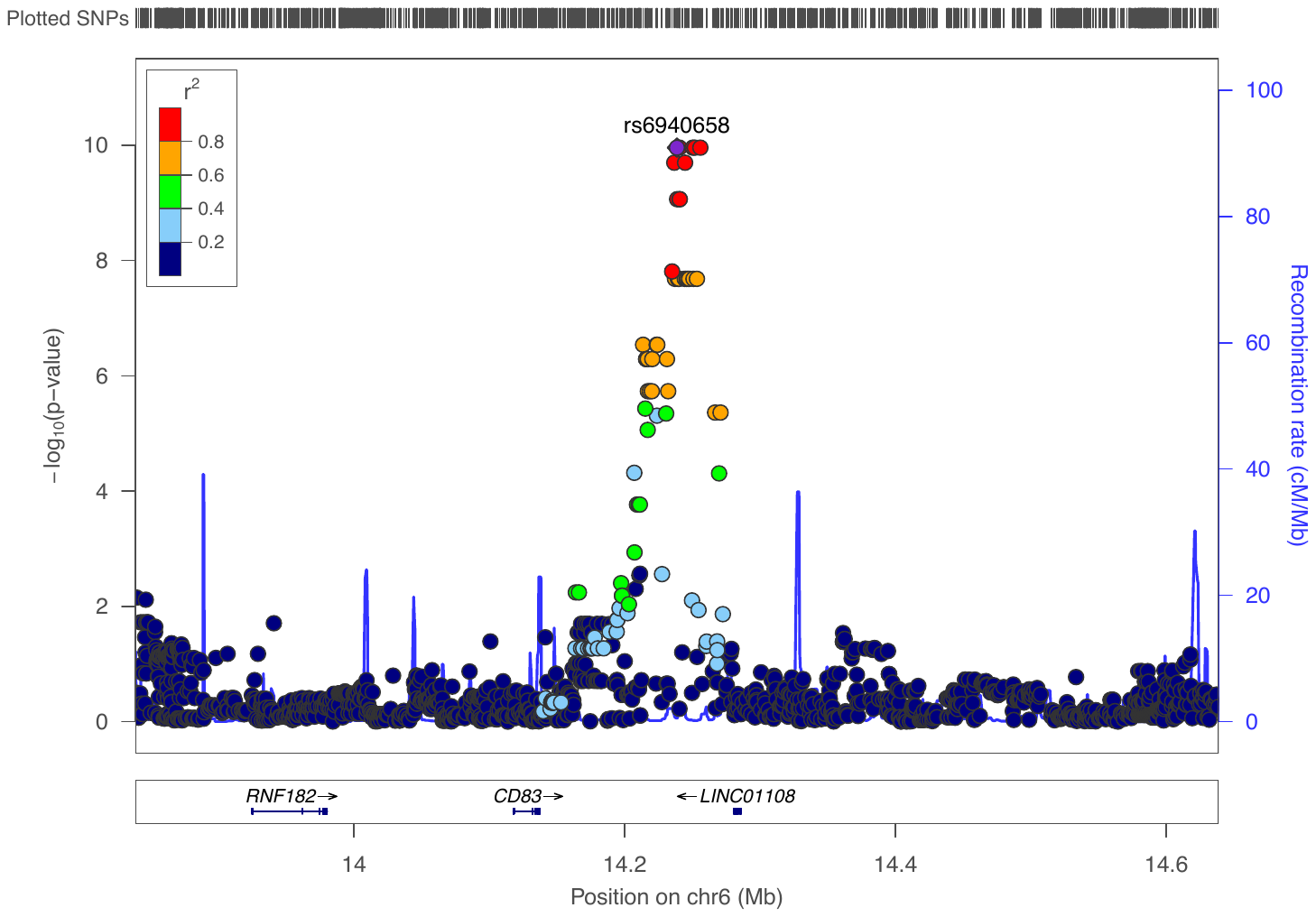

**Figure S5.** LocusZoom plot depicting the GWAS association of CER[N(26)S(19)] to the locus at CD83 on chromosome 6.

The figure depicts the GWAS association of this ceramide species with the locus of the leukocyte surface marker, CD83 in 991 plasma samples. The r^2^ for each SNP is depicted in colour. This finding was supported by three significant genotyped SNPs. The plot was created using the LocusZoom plot tools at http://locuszoom.sph.umich.edu/.

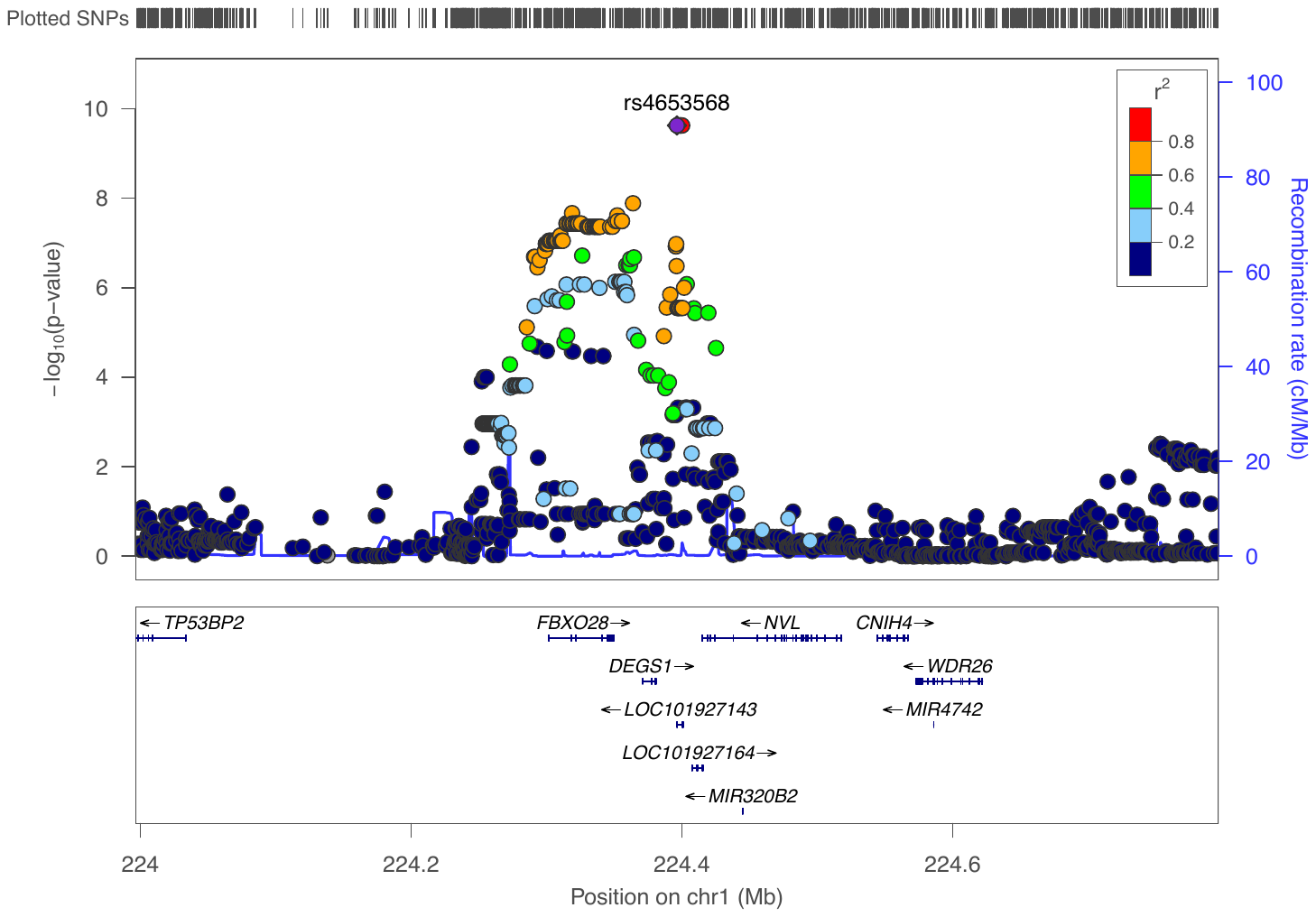

**Figure S6.** LocusZoom plot depicting the GWAS association of the ratio of CER[N(24)S(19)] to its precursor CER[N(24)DS(19)], at the locus of *DEGS1* on chromosome 1.

The figure depicts the GWAS association of this ceramide species with the locus of the desaturase enzyme, DEGS1 in 991 plasma samples. The r^2^ for each SNP is depicted in colour. This finding was supported by two significant genotyped SNPs. The plot was created using the LocusZoom plot tools at http://locuszoom.sph.umich.edu/.

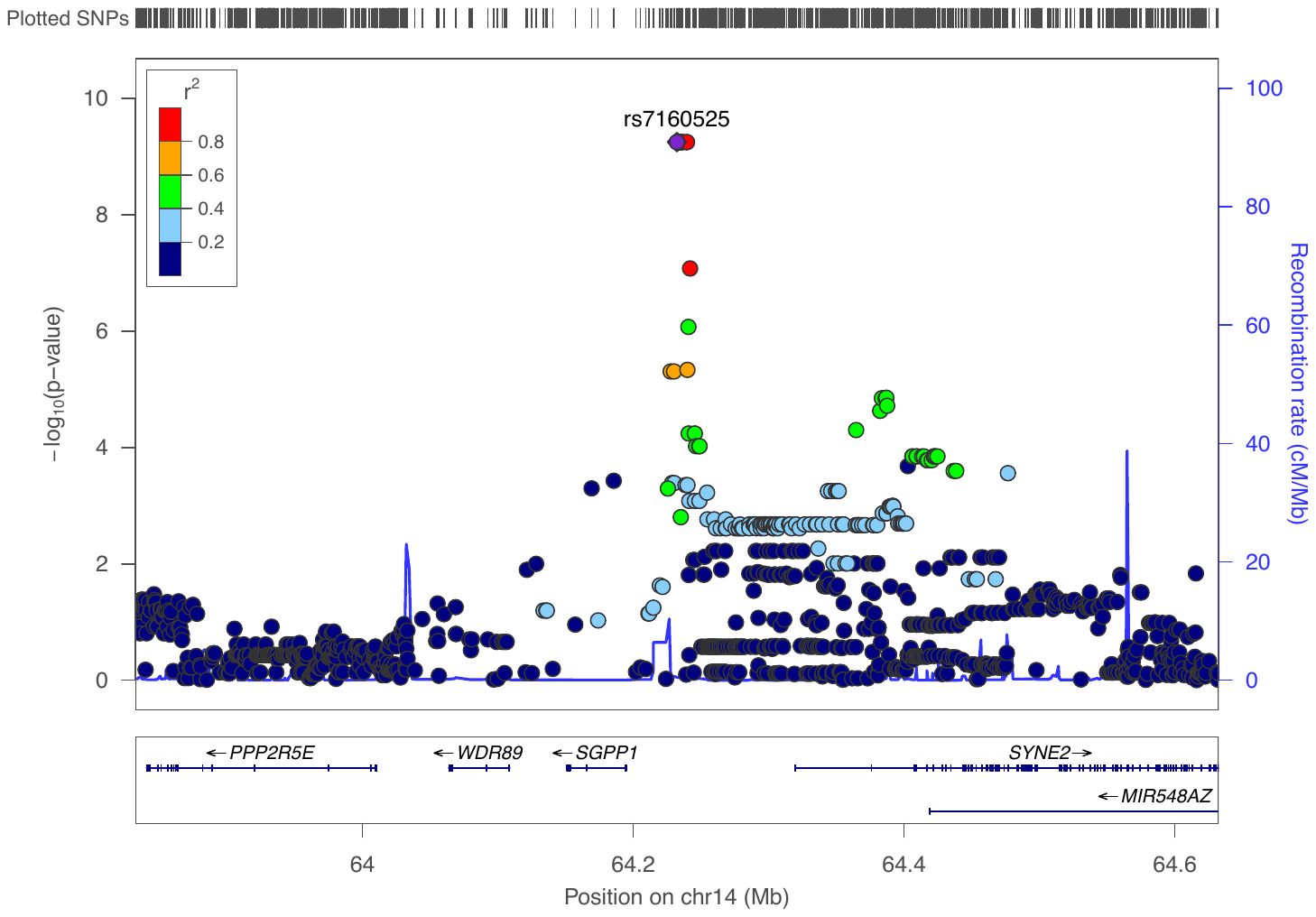

**Figure S7.** LocusZoom plot depicting the GWAS association of CER[N(24)S(16)] to the locus at *SGPP1* on chromosome 14.

The figure depicts the GWAS association of this ceramide species with the locus of the sphingosine 1-phosphate phosphatase enzyme, SGPP1 in 992 plasma samples. The r^2^ for each SNP is depicted in colour. This finding was supported by rs7157785, a GWAS significant genotyped SNP. The plot was created using the LocusZoom plot tools at http://locuszoom.sph.umich.edu/.
